# Environment predicts the maintenance of reproductive isolation in a mosaic hybrid zone of rubber rabbitbrush

**DOI:** 10.1101/2023.08.09.552705

**Authors:** Trevor M. Faske, Alison C. Agneray, Joshua P. Jahner, Carolina Osuna-Mascaró, Lana M. Sheta, Bryce A. Richardson, Elizabeth A. Leger, Thomas L. Parchman

**Affiliations:** Department of Biology, University of Nevada, Reno, NV 89557, USA; Ecology, Evolution, and Conservation Biology, University of Nevada, Reno, NV 89557, USA; Bureau of Land Management, Nevada State Office, Reno, NV 89502, USA; Department of Botany, University of Wyoming, Laramie, Wyoming, 82071, USA; USDA Forest Service, Rocky Mountain Research Station, Moscow, ID 83843, USA

**Author notes:** Corresponding author: Trevor M. Faske, 1664 N. Virginia Street, Department of Biology, 100, University of Nevada, Reno, Reno, NV 89557, USA.

## Abstract

Widely distributed plants of western North America experience divergent selection across environmental gradients, have complex histories shaped by biogeographic barriers and distributional shifts, and often illustrate continuums of reproductive isolation. Rubber rabbitbrush (*Ericameria nauseosa*) is a foundational shrub species that occurs across diverse environments of western North America. Its remarkable phenotypic diversity is currently ascribed to two subspecies – *E. n. nauseosa* and *E. n. consimilis –* and 22 named varieties. We analyzed how genetic variation is partitioned across subspecies, varieties, and environment using high throughput sequencing of reduced representation libraries. We found clear evidence for divergence between the two subspecies, despite largely sympatric distributions. Numerous locations exhibiting admixed ancestry were not geographically localized but were widely distributed across a mosaic hybrid zone. The occurrence of hybrid and subspecific ancestries was strongly predicted by environmental variables as well as the proximity to major ecotones between ecoregions. Although this repeatability illustrates the importance of environmental factors in shaping reproductive isolation, variability in the outcomes of hybridization also indicated these factors likely differ across ecological contexts. There was mixed evidence for the evolutionary cohesiveness of varieties, but several genetically distinct and narrow endemic varieties exhibited admixed subspecific ancestries, hinting at the possibility for transgressive hybridization to contribute to phenotypic novelty and the colonization of new environments in *E. nauseosa*.

## Introduction

Evolutionary biologists have long sought to understand the processes causing phenotypic and genetic divergence between lineages and the origin and maintenance of reproductive isolation (Nosil & Feder, 2012; Schluter, 2001; Thompson, 2005). As a result, much work has focused on populations spanning the continuum of divergence and reproductive isolation that may characterize the progression of speciation (Bolnick et al., 2023; Stankowski & Ravinet, 2021). Widely distributed plant taxa of western North America often display extensive variation in phenotype and connectivity due to divergent selection across heterogeneous landscapes that support environmental gradients (e.g., Baughman et al., 2019), as well as historical periods of isolation owing to biogeographical barriers and distributional shifts associated with Quaternary climate oscillations (Dynesius & Jansson, 2000; Massatti et al., 2018; Massatti & Knowles, 2020; Shafer et al., 2010). Temporal and spatial variation in these processes may lead to a continuum of phenotypic and genetic divergence with variable reproductive isolation among populations and species (i.e., syngameons; Buck et al., 2023; Chhatre et al., 2018; Grant, 1981; Lotsy, 1925). Plant groups composed of complexes of interbreeding and isolated lineages are relatively common in the American West due to topographic and environmental complexity, and can be valuable for understanding how ecological variation underlies adaptation, speciation, and hybridization.

The outcomes of hybridization between lineages depend on hybrid fitness and the genetic basis of reproductive isolation (Barton, 2001). Understanding the factors that prevent or promote hybridization can provide insight into the genetic and environmental causes of divergence, isolation, and gene flow. Ecological factors, such as variation in specialization, species interactions, or phenology, can mediate prezygotic barriers to reproduction (e.g., Hood et al., 2019). Postzygotic barriers can also be influenced by ecological factors, such as reduced hybrid fitness in either parental environment, or by purely genetic interactions (Maheshwari & Barbash, 2011; Orr, 1996). Gene flow occurring through hybridization can obscure species boundaries, blur phylogeny, and contribute to taxonomic uncertainty (McVay et al., 2015; Zhang et al., 2021). Further, hybridization can contribute to adaptation and evolutionary novelty through both adaptive introgression (e.g., Calfee et al., 2020; Giska et al., 2019) and transgressive variation in hybrids that may facilitate the colonization of novel environments (Gompert et al., 2006; Hodel et al., 2022; Rieseberg et al., 1999). While many instances of hybridization are structured as geographic clines across major ecotones (Barton & Hewitt, 1985; Endler, 1977), more complex outcomes can arise in the form of mosaic hybrid zones, where hybrids between lineages occur at multiple locations across a mosaic of environmental variation (Rand & Harrison, 1989). Geography and environment are decoupled in mosaic hybrid zones, offering replicate views into the predictability of reproductive isolation and the environmental factors underlying it. Here we evaluate the ecological predictors of divergence and secondary contact in a morphologically and taxonomically diverse foundational shrub species found throughout the geologically complex, arid regions of western North America.

*Ericameria* (Asteraceae) is a genus of approximately 70 shrub species distributed in western North America. Rubber rabbitbrush (*Ericameria nauseosa*) is broadly distributed and abundant across desert, woodland, and arid montane habitats, where it thrives as a primary successional colonizer of arid and disturbed sites. It is a foundational species with deep root systems that stabilize soil for arid plant communities (Donovan & Ehleringer, 1994) and provides important resources for birds, mammals, and insects. It hosts an exceptional number of gall-forming insects (Fernandes et al., 2000; Floate et al., 1996; Mcarthur et al., 1986) and represents a keystone late-season flowering resource for diverse pollinator communities (Carril et al., 2018; Hansen, 1986; McArthur et al., 1979; Ogle et al., 2007; Rogers, 1979). Across the environments and geographic regions it occupies, *E. nauseosa* displays substantial phenotypic variation in its stature, morphology, and coloration, and is noted for its exceptional phytochemical diversity (Hegerhorst et al., 1987a,b; Weber et al., 1985). Pronounced population differentiation in germination timing has also illustrated local adaptation to variation in temperature across its distribution (Meyer et al., 1989). Given its ecological significance, phenotypic and genetic variation in *E. nauseosa* are likely to have extended community and ecosystem level consequences (Grady et al., 2011; Whitham et al., 2003) and to have value for restoration of native plant communities across much of western North America (Faske et al., 2021).

Due to extensive phenotypic differentiation, taxonomists have recognized numerous subspecies and varieties within *E. nauseosa*, and taxonomic revisions have been common. *Ericameria nauseosa* was moved from *Chrysothamnus* to *Ericameria* (Nesom & Baird, 1993) and subspecific classifications have shifted from 22 recognized subspecies (Anderson, 1986a; Hall, 1919) to the current designation of two recognized subspecies and 22 named varieties (Nesom & Baird, 1993). The two recognized subspecies (*E. n. nauseosa* [grey] and *E. n. consimilis* [green]) have overlapping distributions but are phenotypically distinguishable by variation in grey and green pigmentation in the leaves and stems (Figure 1A; Anderson, 1986b). The 22 named varieties vary in vegetative and floral morphology and in distribution, with some varieties spanning much of western North America and others being narrowly endemic soil specialists (Anderson, 1986a; Nesom & Baird, 1993). There is distinctive phytochemical variation between both subspecies and among varieties (Hegerhorst et al., 1987a,b; Weber et al., 1985) and this variation is associated with distinct insect and herbivore communities (Floate et al., 1996; Mcarthur et al., 1986; McArthur et al., 1979; Wangberg, 1981). The subspecies and several varieties are sympatric in some locales, and while hybridization has been noted or suspected (Anderson, 1973; Hanks et al., 1975), the persistence of differentiation suggests some degree of reproductive isolation (Anderson, 1986b; Anderson & Reveal, 1966; McArthur et al., 1979). Thus, the phenotypic and taxonomic diversity in *E. nauseosa* could reflect diversification driven by adaptation to different environments that has resulted in a series of independently evolving lineages across a continuum of diversification.

**Figure 1:**
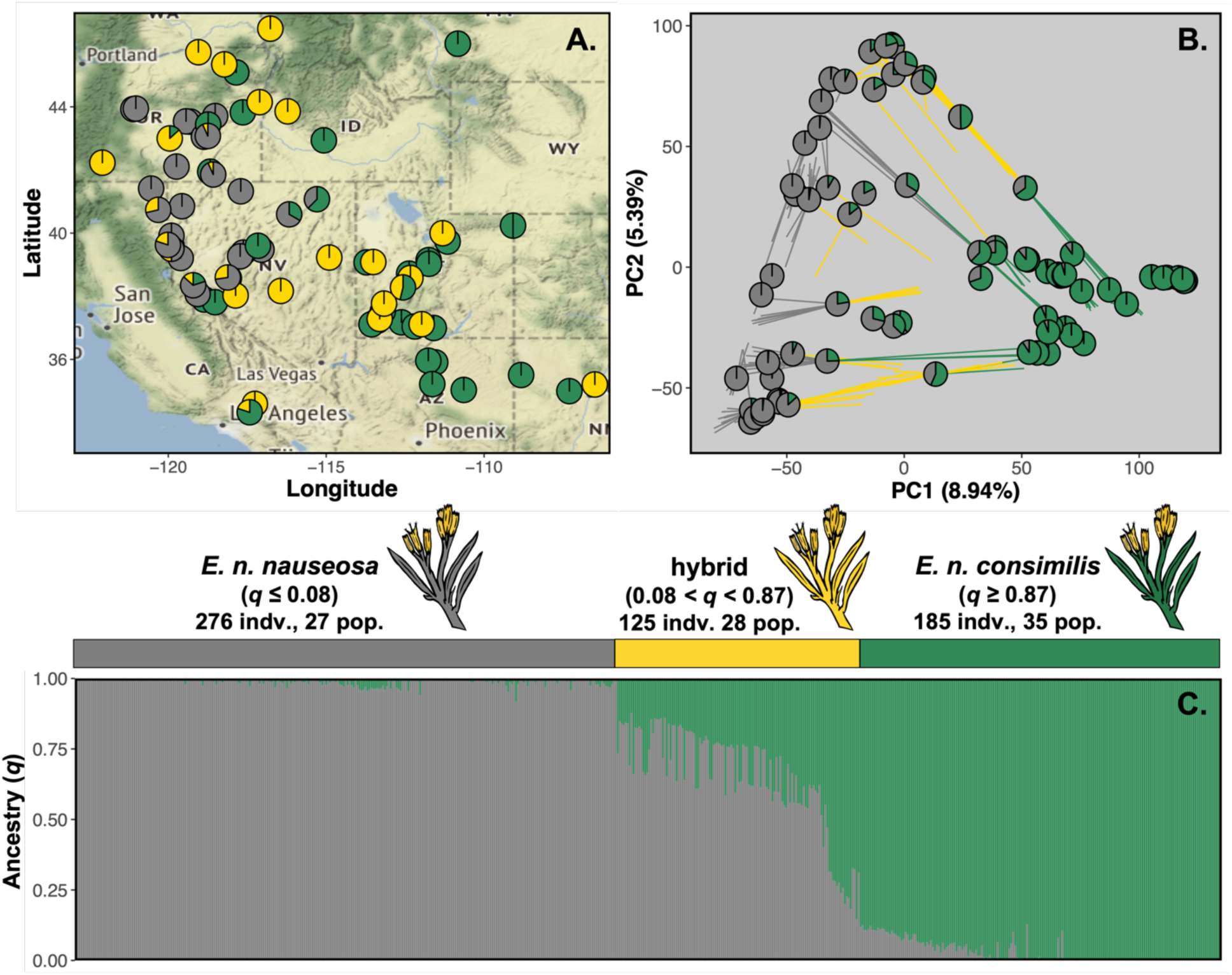
*E. nauseosa* lineages, *E. n. nauseosa* and *E. n. consimilis,* exhibit genetic differentiation and admixture occurs in multiple locations. (**A**) Map corresponding to the sampling locations. Pie chart colors correspond to the proportion of individuals at a locality assigned to each lineage or as a hybrid. (**B**) Principal component analysis (PCA) illustrates genetic differentiation between lineages with the presence of hybrid individuals. Circles represent mean PC values for populations, pie charts reflect ancestry, and lines connect PC values for individuals to population means. Yellow lines represent hybrid individuals with ancestry coefficients 0.08 < q < 0.87. (**C**) Ancestry coefficients (*q*) from the hierarchical Bayesian model of entropy of *k* = 2 for each individual, ordered by PC1. Color proportion of each bar ancestry proportion. See Figure S1 for similar analyses conducted on a downsampled dataset.

As the few prior population genetic analyses of the group have been limited to single subspecies or varieties (Faske et al., 2021; Gang & Weber, 1995), we lack an understanding of the evolutionary cohesiveness of taxonomic designations and the potential for genetic differentiation and reproductive isolation among such groups. Furthermore, while hybridization among subspecies and varieties has been anecdotally suspected, we have no empirical understanding of its actual extent. High-throughput sequencing approaches have improved our ability to resolve the early phases of diversification (Buck et al., 2023; Chhatre et al., 2018; R. Fu et al., 2022; Wagner et al., 2013), to characterize the influence of environment on genetic variation (Forester et al., 2016, 2018; Rellstab et al., 2015), and quantify ancestry variation among and within admixed populations (Gompert et al., 2017; Moran et al., 2021). Geographic variation in the distribution of hybrid ancestry can provide insights into the ecological and environmental factors mediating reproductive isolation and shaping divergence in diverse plant assemblages.

Here we quantified range-wide genetic variation in *E. nauseosa* using high-throughput reduced representation sequencing for 76 sampling localities spanning taxonomic variation and a substantial portion of the geographic and environmental breadth of its distribution. We first characterized the degree of genetic structure between and within subspecies, and asked how it has been shaped by geography, environment, and history. Second, given past observations of suspected or confirmed hybrids, we quantified the extent and predictors of hybridization across genetic lineages. Finally, given that the broad morphological and ecotypic diversity in *E. nauseosa* has been circumscribed into 22 named varieties, we evaluated the evolutionary and genetic cohesiveness of putative varietal designations. Our results reveal a continuum of genetic divergence and reproductive isolation spanning lineages, environments, and phenotypes that will guide future population genomic work in this syngameon-like foundational species and provide useful perspective for restoration in the Intermountain West.

## Methods

### Plant sampling and taxonomic identification

We collected fresh leaf tissue from 76 sampling locations (608 individuals; 2 - 15 per location; Table S1), spanning a substantial portion of the *E. nauseosa* range in the western United States (Figure 1). During field collection, plants were identified to subspecies and variety for most locations (Cronquist, 1994). Our sampling ultimately captured 14 of the 22 named varieties. Current subspecific designations of the sampled varieties follows *E. n. nauseosa* (varieties *bigelovii*, *graveolens*, *hololeuca*, *iridis*, *latisquamea*, *nana*, *salicifolia*, *speciosa*) and *E. n. consimilis* (varieties *arenaria*, *juncea*, *mohavensis*, *nitida*, *oreophila*, *turbinata*). Individuals from locations where phenotypic assignment to variety proved difficult were designated as unknown; this was at times due to the potential existence of more than a single subspecies or variety in a single location and the difficulty of using phenotypic characters in dichotomous keys. Three samples of *E. discoidia,* a sister taxon known to hybridize with *E. nauseosa* (Anderson & Reveal, 1966; Roberts & Urbatsch, 2003), were additionally included for phylogenetic outgroup purposes.

### Reduced representation sequencing, bioinformatic processing, and variant calling

Dried leaf material from each plant was ground into powder form using a Qiagen TissueLyser and DNA was extracted using Qiagen DNeasy Plant kits (Qiagen, Valencia CA). Reduced representation libraries were prepared using a double-digest restriction-site associated DNA sequencing (ddRADseq) method (Parchman et al., 2012; Peterson et al., 2012). The restriction endonucleases *Eco*RI and *Mse*I were used to digest genomic DNA and uniquely barcoded Illumina adaptors were subsequently ligated to the resulting fragments using T4 DNA ligase (New England Biolabs, Ipswich MA). Barcoded fragments were PCR amplified with Iproof DNA polymerase (BioRad, Hercules CA), and fragments ranging from 350 to 450 bp were size selected using a Pippin Prep quantitative electrophoresis unit (Sage Science, Inc). Single-end sequencing (100 bp read lengths) was performed using two lanes on an Illumina HiSeq 4000 platform at the University of Wisconsin.

Sequences representing potential contaminants (*E. coli* and PhiX) or various Illumina associated oligos were detected and discarded using a pipeline of Perl and bash scripts (http://github.com/ncgr/tapioca). A custom Perl script was then used to correct barcode sequencing errors, to trim cut site and barcoded oligo associated bases, and to demultiplex reads by individual. As a reference genome is not available for *E. nauseosa* or a close relative, we used a *de novo* assembly of unique reads to create a consensus reference of genomic regions sampled by our reduced representation approach (reference assembly, hereafter). Parameters for this step were optimized using shell scripts and documentation provided for dDocent (Puritz et al., 2014) (cutoffs: individual = 8, coverage = 5; clustering similarity: -c .92), and the reference assembly was produced using CD-HIT-EST V4.8.1 (Fu et al., 2012, Li & Godzik 2006). Demultiplexed reads for each individual were mapped to the reference assembly using BWA-MEM V0.7.17 (Li, 2013). Sequence variants were identified using SAMTOOLS V1.9 and BCFTOOLS V1.9 (Li et al., 2009), and subsequent filtering was completed using VCFTOOLS V0.1.16 (Danecek et al., 2011). We retained only biallelic loci covered by reads present in at least 70% of individuals, thinned to one locus per contig to reduce effects of linkage disequilibrium and sequencing error. Additionally, individuals missing data for greater than 40% of loci were removed before further analyses. After additional filtering with custom Python scripts, we retained loci with minor allele frequency (MAF) ≥ 0.02, mean depth across individuals ≥ 3 or < 25, and alternate allele call quality ≥ 750. We removed loci with excessive coverage and only retained loci with two alleles present to ameliorate genotyping bias from the potential misassembly of paralogous genomic regions (Hapke & Thiele, 2016; McKinney et al., 2018). Lastly, we retained loci with *F*_IS_ > –0.5, as misassembly of paralogous genomic regions can lead to abnormal heterozygosity (Hohenlohe et al., 2013; McKinney et al., 2017). To detect duplicated, clonal, or highly related individuals, we quantified relatedness among all pairs of individuals within each population using VCFTOOLS, employing the cutoff criteria provided by the KING method (Manichaikul et al., 2010; Turner et al., 2022).

### Landscape genetic structure

To estimate genotype probabilities for each individual at each locus, infer the number of ancestral genetic clusters (*k*), and estimate individual ancestry coefficients (*q*), we utilized a hierarchical Bayesian model that incorporates genotype uncertainty (ENTROPY V2.0; Gompert et al., 2014; Shastry et al., 2021). The model employed by ENTROPY utilizes an allele frequency prior and incorporates genotype uncertainty arising from sequencing and alignment error, as well as stochastic variation among individuals and loci in sequencing depth during parameter estimation (Gompert et al., 2014). Using genotype likelihoods calculated from SAMTOOLS, we executed *k*-means clustering of the first five PCs to generate starting values of ancestry coefficients (*q*) to seed the MCMC (Gompert et al., 2014). ENTROPY was used to estimate genotype probabilities and ancestry coefficients for models based on a range of ancestral genetic clusters (*k*). We ran 60,000 MCMC iterations across 4 chains with a burn-in of 10,000, thinning every tenth step, for models based on each level of *k*. Genotype probabilities from the top five models based on DIC (*k* = 2 – 6) were averaged across all chains and used for all subsequent population genetic analyses (Table S2). To verify that ordination and ancestry analyses were not biased by uneven sampling (McVean, 2009; Novembre & Stephens, 2008; Sinclair & Hobbs, 2009), we filtered variants, inferred ancestry coefficients, and conducted population genetic analyses as described below on a reduced data set with even sampling across sampling locations and subspecies (231 individuals, 47 locations). Because patterns of clustering and population differentiation were qualitatively similar for both datasets (see Figure 1, Supplementary Figure 1), irrespective of sampling density, we used the full data set for analyses detailed below. As a complementary model-free approach to quantify genetic variation among all individuals spanning the named varieties and subspecies, we conducted a principal component analysis (PCA) on genotype probabilities, following standardization proposed by Patterson et al., (2006).

Both PCA and ENTROPY suggested the two lineages were genetically differentiated and sympatric (Figure 1), with hybrids throughout the sampled range, so we additionally conducted population genomic analyses separately for sets of individuals with pure ancestry for each lineage (excluding hybrids). We use the term ‘lineage’ along with the subspecific nomenclature (*E. n. nauseosa* and *E. n. consimilis*) to refer to these genetic groups, overriding field identifications in cases where subspecies designation was incongruent with genetic data. Specifically, individuals were categorized by ancestry coefficients (*q*) based on cut-offs that made sense geographically and biologically (Figure 1C; *E. n. nauseosa* [grey] < 0.08 *≤* hybrid *≤* 0.87 < *E. n. consimilis* [green]). Sampling locations containing individuals across multiple ancestral categorizations (i.e., *E. n. nauseosa*; *E. n. consimilis*; or hybrid ancestry *n* = 13 locations) were treated as separate populations. Variants were sampled independently from the genotype probabilities and re-filtered with a MAF ≥ 0.02 (*E. n. nauseosa*: 18,366 loci, 276 indv., 27 pop.; *E. n. consimilis*: 15,618 loci, 185 indv., 34 pop.).

We visualized population structure within each lineage (*E. n. nauseosa* and *E. n. consimilis*) using PCA. To evaluate the concordance of population genetic variation and geography, we applied Procrustes rotation of the first two PC axes onto population latitude and longitude coordinates (Wang et al., 2010). To further quantify the spatial scale of population differentiation, we used Random Forest models to assess the degree to which individuals could be classified to their population of origin based on genotypic data alone. We conducted this analysis separately within *E. n. nauseosa* and *E. n. consimilis* and, again, for the full set of individuals. Sampling location was used as the response variable and principal component scores for each individual from PCAs of genotype probabilities were treated as predictors. The number of PC axes was selected using Tracy-Widom tests in R (R Core Team, 2020) package LEA (Gain & François, 2021) with an ɑ ≤ 0.001 (overall = 59 PCs; *E. n. nauseosa* = 25 PCs, *E. n. consimilis* = 27 PCs). Models were analyzed using the R package RANDOM FOREST (Breiman, 2001), and accuracy was assessed using the out-of-bag error (OOB). As a metric of genetic differentiation among populations within and between each lineage, we estimated with multilocus *F*_ST_ calculated with the HIERFSTAT R package (Goudet & Jombart, 2020). We used Mantel tests (Mantel, 1967) to quantify isolation-by-distance (IBD; Wright, 1943) by associating genetic distances (Nei’s *D*; Nei, 1972) with geographic distances within and between each lineage. Geographic distances among populations were calculated as haversine distance using the GEOSPHERE (Hijmans, 2017) R package, and the Procrustes rotation and Mantel tests were calculated using the VEGAN (Oksanen et al., 2019) R package.

Because variation in genetic diversity parameters could provide cursory information about historical demography of the subspecific lineages, we estimated a suite of population level metrics. Specifically, we estimated nucleotide diversity (*θ_π_*; Nei & Li, 1979), the Watterson estimator (*θ_W_*; Watterson, 1975) and Tajima’s *D* (Tajima, 1989) for each population (min. 3 individuals) using methods implemented in ANGSD V 0.923 that incorporate genotype uncertainty (Korneliussen et al., 2013, 2014). Additionally, individual inbreeding coefficients (*F*) were calculated while incorporating genotype uncertainty using NGSF (Vieira et al., 2016), as implemented in the ANGSD-WRAPPER V 0.933 (Durvasula et al., 2016; see Supplementary methods).

### Environmental and ecological influences of lineage differentiation and hybridization

Results from ENTROPY and PCA revealed that individuals of pure *E. n. nauseosa* and *E. n. consimilis* ancestry co-occurred in some locations, and many locations exhibiting hybrid ancestry were widely dispersed across the sampled region. Thus, to consider the extent to which extrinsic factors might shape reproductive isolation, or the lack thereof, we analyzed the extent to which environmental variation predicted the spatial distribution of lineage and hybrid ancestry. We characterized environmental variation across sampling sites with 30 climate variables of the Climatic Water Deficit Toolbox (Dilts et al., 2015). This dataset provides estimates of potential evapotranspiration, actual evapotranspiration, climatic water deficit, and 30-year PRISM normals (1981 – 2010) at 500m × 500m resolution and has been shown to predict aspects of spatial and distributional variation across plant communities (Lutz et al., 2010; Stephenson, 1998). Variables were reduced from 30 to 10 (based on Pearson’s |*r*| < 0.75) to reduce the potential influence of multicollinearity. Full descriptions of these variables and their values for each sample location can be found in Supplementary Tables 3 and 4.

As both ecological and environmental factors can influence differentiation between lineages and reproductive isolation after secondary contact, we explored multiple facets of genotype-environment associations. To interpret environmental associations appropriately, the confounding correlation of geography and environment (Supplementary Figure 2) was taken into consideration throughout the following analyses. We first quantified the association between genetic and environmental distances, or isolation-by-environment (IBE; Wang & Bradburd, 2014) within and between each lineage using Mantel tests. We then estimated the magnitude and directionality of each environmental variable’s influence on spatial genetic structure through redundancy analyses (RDA). We conducted three additional analyses aiming to: (1) decouple the combined and independent influences of environment and geography on lineage divergence (i.e., ancestry coefficients) using a variance partitioning approach; (2) assess the predictability of lineage and hybrid assignment from environmental data using Random Forest; and (3) examine the association of individual environmental and geographic predictors with both *q* and lineage assignment of each population. Finally, as hybridization may occur more commonly across ecotones, we measured the average minimum geographic distance to ecoregion boundary between the sampling locations within each lineage and hybrids based on the EPA guided North American ecoregion level 2 (Omernik & Griffith, 2014; see Supplementary Methods).

### Assessing the evolutionary cohesiveness of varieties

We inferred phylogenetic relationships spanning locations covering the two lineages and hybrids, as well as 14 of the named varieties. We included samples of *E. discoidea* for outgroup purposes. We used this analysis as an additional measure of divergence between the two lineages, and to evaluate the evolutionary cohesiveness of the putative varieties. For this analysis, we used field-designated subspecies and varieties and did not adjust them based on previous results. To ease computational investment and facilitate clear graphical visualization, we subsetted the data by randomly selecting up to six individuals from a reduced number of locations (231 individuals across 47 populations with even sampling [4-6 inds] across locations and subspecies). As the phylogenetic method used does not account for admixture, an additional tree was generated excluding hybrid individuals to complement inference. Variants were refiltered using a MAF ≥ 0.02 cutoff and resulting vcf files were converted into PHYLIP format using VCF2PHYLIP V 2.0 (Ortiz, 2019). We then reconstructed a Maximum-Likelihood phylogenetic tree using IQ-TREE V 2.0.3 (Minh et al., 2020; Nguyen et al., 2015) with the PHYLIP file as input. We used the “Model Finder Plus” parameter (-m MFP) to determine the model that minimized the BIC score, implemented the ascertainment bias correction method (ASC; Lewis, 2001), and applied the ultrafast bootstrap option with 1000 bootstrap replicates (-bb 1000) to measure branch support. We visualized and annotated the resulting tree using GGTREE (Yu et al., 2017) in R. Additionally, to assess degree of range sympatry among the lineages and varieties, range maps from all sampled varieties were digitized from Anderson (1986b) with area and pairwise percent overlap calculated. Digitizing and georeferencing was performed manually in Inkscape V 0.92.5 (Inkscape project 2020) with shapefile generation, area calculations, and graphics were created using the R packages RASTER, RGEOS, SP, and SF (Bivand & Rundel 2020; Hijmans 2022; Pebesma, 2018; Pebesma & Bivand 2005).

## Results

### Landscape genetic structure

After contaminant cleaning, mapping, variant calling, and filtering, we retained 22,917 loci in 586 individuals spanning 76 sampling locations (2 – 15 inds. per locale). There was no evidence of highly related individuals in our sampled locations (mean kinship coefficient = 0.039; only one pairing above the full-sibling estimate), consistent with outcrossing, and suggesting that subsequent population genetic analyses were not confounded by the presence of clones or familial relationships. Genetic differentiation between two lineages largely corresponding with subspecific designations (*E. n. nauseosa* and *E. n consimilis*) was evident in both the PCA and ancestry coefficients for the *k* = 2 ENTROPY model (Figure 1B, C). The first PC axis (*PVE =* 8.94%) partitioned the two lineages and was highly correlated to ancestry (|*r|* = 0.968), while the second PC axis (*PVE =* 5.39%) was associated with population structure within each lineage (i.e., geography; PC2 – latitude: |*r|* = 0.425, PC2 – longitude: |*r| =* 0.350). Thus, unless specified otherwise, for subsequent analyses we categorized individuals based on ancestry coefficients as hybrid or belonging to one of the two lineages (*E. n. nauseosa*: *q* ≤ 0.08; *E. n. consimilis* : *q* ≥ 0.87; hybrid: 0.08 < *q* < 0.87, 125 indv., 28 pop.). Despite substantial geographic overlap (Figure 1A), the two lineages exhibited clear genetic differentiation (*F*_ST_ = 0.111, 95% bootstrap CI = 0.107 – 0.114).

When the two lineages were analyzed separately, population structure within each was strongly predicted by geographic variation and was often clear at fine spatial scales (<30 km, Figure 2). Individuals from the same populations generally formed tight clusters in PCA space and Random Forest analyses correctly categorized individuals to their sampling origin at extremely high rates. Specifically, individual assignment accuracy was 94.37% with 69 / 76 sampling locations having at most one individual incorrectly classified for the full set of samples. Individual assignment accuracy within each lineage was similarly high (95.29% and 92.43%), with 26 / 27 (*E. n. nauseosa*) and 31 / 35 (*E. n. consimilis*) sampling locations having at most one individual incorrectly classified to location. Procrustes rotation correlations of geography onto the first 2 PCs was high, indicating that genetic structure was strongly predicted by geography (*E. n. nauseosa*: Procrustes *r* = 0. 761; *E. n. consimlis*: Procrustes *r* = 0.572). Additionally, Mantel tests of geographic distance and genetic distance (IBD) within lineages were strong (*E. n. nauseosa*: Mantel *r* = 0. 406, *P* = 0.001; *E. n. consimilis*: Mantel *r* = 0.239, *P* = 0.006) but absent when IBD was assessed comparing populations spanning the two lineages (Mantel *r* = −0.083, *P* = 0.790; Figure 2C); consistent with geographically overlapping distributions of two differentiated lineages (see Supplementary Figure 2 for full associations). Across the lineages and hybrids, genetic diversity estimates were high and inbreeding coefficients (*F*) were low. *Ericameria n. nauseosa* had the greatest genetic diversity and lowest inbreeding, on average, with the hybrids having intermediate values between *E. n. nauseosa* and *E. n. consimilis* (Supplementary Figure 3). All population estimates of Tajima’s *D* were negative, but Tajima’s *D* was associated with latitude in opposite directions in *E. n. nauseosa* and *E. n. consimilis* (Figure 2D).

**Figure 2:**
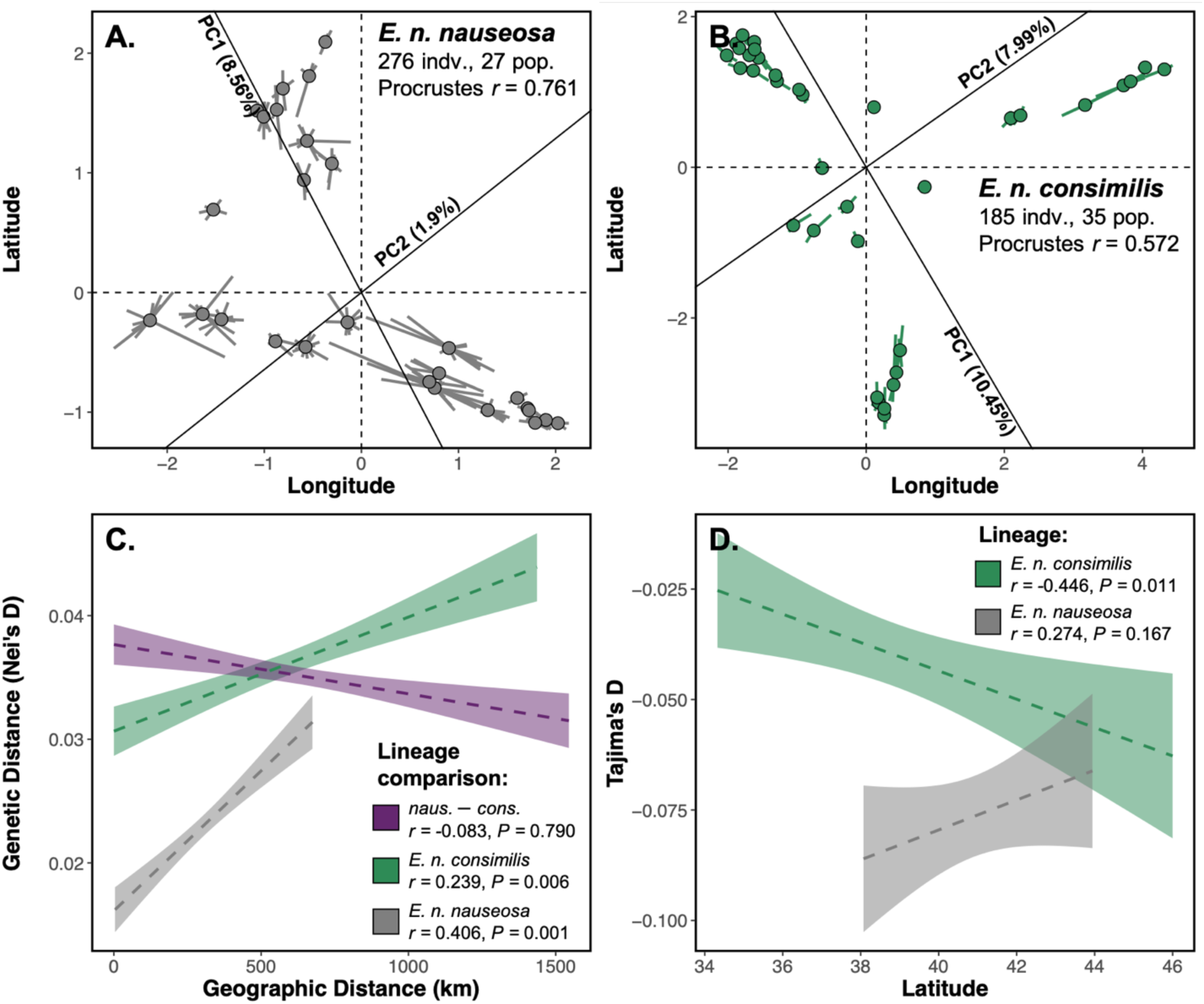
*E. n nauseosa* and *E. n. consimilis* form independently evolving lineages, exhibiting strong population structure, patterns of IBD, and variation in demographic history. Population genetic structure within both (A) *E. n. nauseosa* and (B) *E. n. consimilis* is demonstrated using a Procrustes rotation of the first 2 PC axes onto latitude/longitude. Circles represent population means connected to lines representing individual PC scores from each population. (**C**) Strong patterns of isolation-by-distance (IBD) were present within the two lineages but not when comparing populations among and within both lineages (shown in purple). (**D**) Tajima’s *D* is negative in both lineages and associated with latitude, but in opposite directions. These results potentially highlight differences in post-glaciation expansion routes between the two lineages.

### Environmental and ecological influences of lineage differentiation and hybridization

Multiple analyses indicated the influence of both environment and geography on spatial patterns of genetic variation. Isolation-by-environment was weakly evident within each lineage (*E. n. nauseosa*: Mantel *r* = 0. 191, *P* = 0.054; *E. n. consimilis*: Mantel *r* = 0.150, *P* = 0.097) but not when analyzed across both lineages (Mantel *r* = −0.048 *P* = 0.677; Figures 3A, Supplementary Figure 2), as expected given evidence for genetic divergence among lineages despite substantially overlapping distributions. Geographic and environmental variables explained genetic variation between the two lineages and were heavily loaded on the first RDA axis (*PVE* = 52.31%; Figure 3B). Geographic locality was unsurprisingly among the largest contributors to genetic differentiation between the lineages with increasing longitude (i.e., the more eastern part of the range) associated with the distribution of *E. n. consimilis* ancestry. Environmental variables also explained ancestry variation with increased cumulative and fall actual evapotranspiration (CumlAET and FallAET), heatload, and annual mean temperature (AnnTemp) associated with *E. n. consimilis* ancestry, and increased values of slope and precipitation seasonality (PrcpSeas) associated with *E. n. nauseosa* ancestry. Individuals with hybrid ancestry were associated with intermediate values of these variables (Fig. 3B). Additionally, the second RDA axis splits the hybrids into two clusters with one associated with increasing soil water capacity (SoilAWC) and decreasing elevation and slope (Figure 3B).

**Figure 3:**
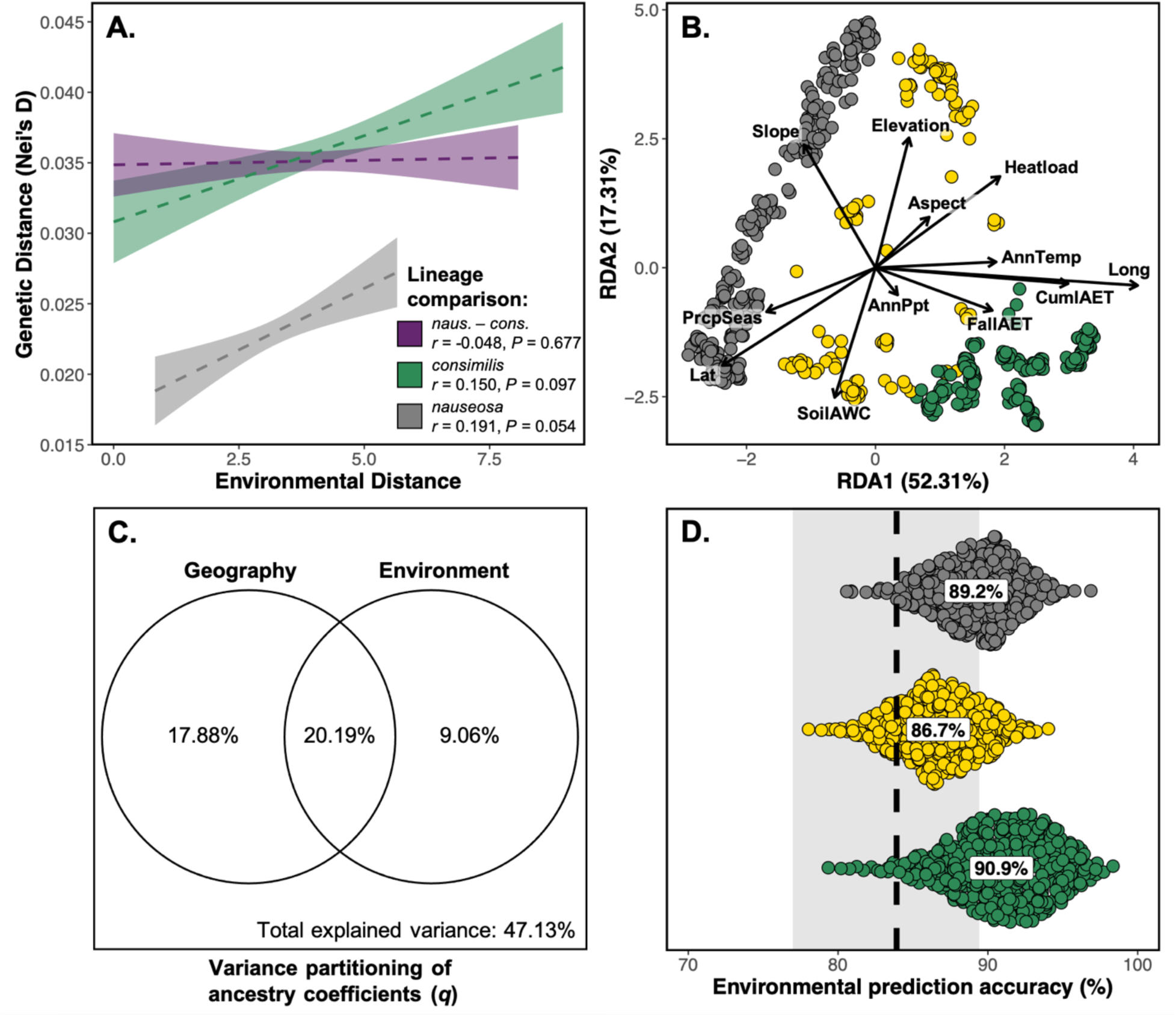
Environmental variation predicts differentiation and occurrence of hybridization between *E. n. nauseosa* and *E. n. consimilis* lineages. (**A**) Strong patterns of isolation-by-environment (IBE) were present within the two lineages but not when comparing populations among and within both lineages (shown in purple). (**B**) Environmental variation associated with genetic structure between the lineages illustrated by RDA. Axes were rotated in order to match that of the PCA in Figure 1 and environmental loadings were scaled (4.36×) for graphical representation. (**C**) Venn diagram illustrating partial variance of environment and geographic effects on ancestry coefficients in terms of adjusted *r*^2^. (**D**) Environment alone predicts an individuals’ classification to either lineage or as a hybrid using Random Forest. Points represent permuted accuracy within each lineage with means for each shown in text. Dashed line and shaded bar represent the overall mean predictive accuracy (83.9%) and 95% confidence interval.

In variance partitioning analyses, environment and geography explained a combined 47.1% of the total variance in *q*, but when taken individually, environment explained 29.3% and geography explained 38.1%. When surveying the partial contributions of environment and geography independent of correlative effects of each other, the combined partial effects contributed the most (adj. *r*^2^ = 20.2%), followed by geography (adj. *r*^2^ = 0.18), and then environment (adj. *r*^2^ = 0.09; Figure 3C). Environmental variation was strongly predictive of ancestry in Random Forest models, which had an overall accuracy of 83.9% (range: 77.0 – 89.4%) in predicting an individual’s lineage classification (mean *r*^2^ = 0.860, RMSE = 0.160; Figure 4). Classification accuracies for *E. n. nauseosa* and *E. n. consimilis* were 89.2% and 90.9%, respectively, while the classification accuracy for hybrids was 86.7% (Figure 3D). Cumulative AET was overwhelmingly the strongest predictor of *q*, followed by annual precipitation, slope, and FallAET.

**Figure 4:**
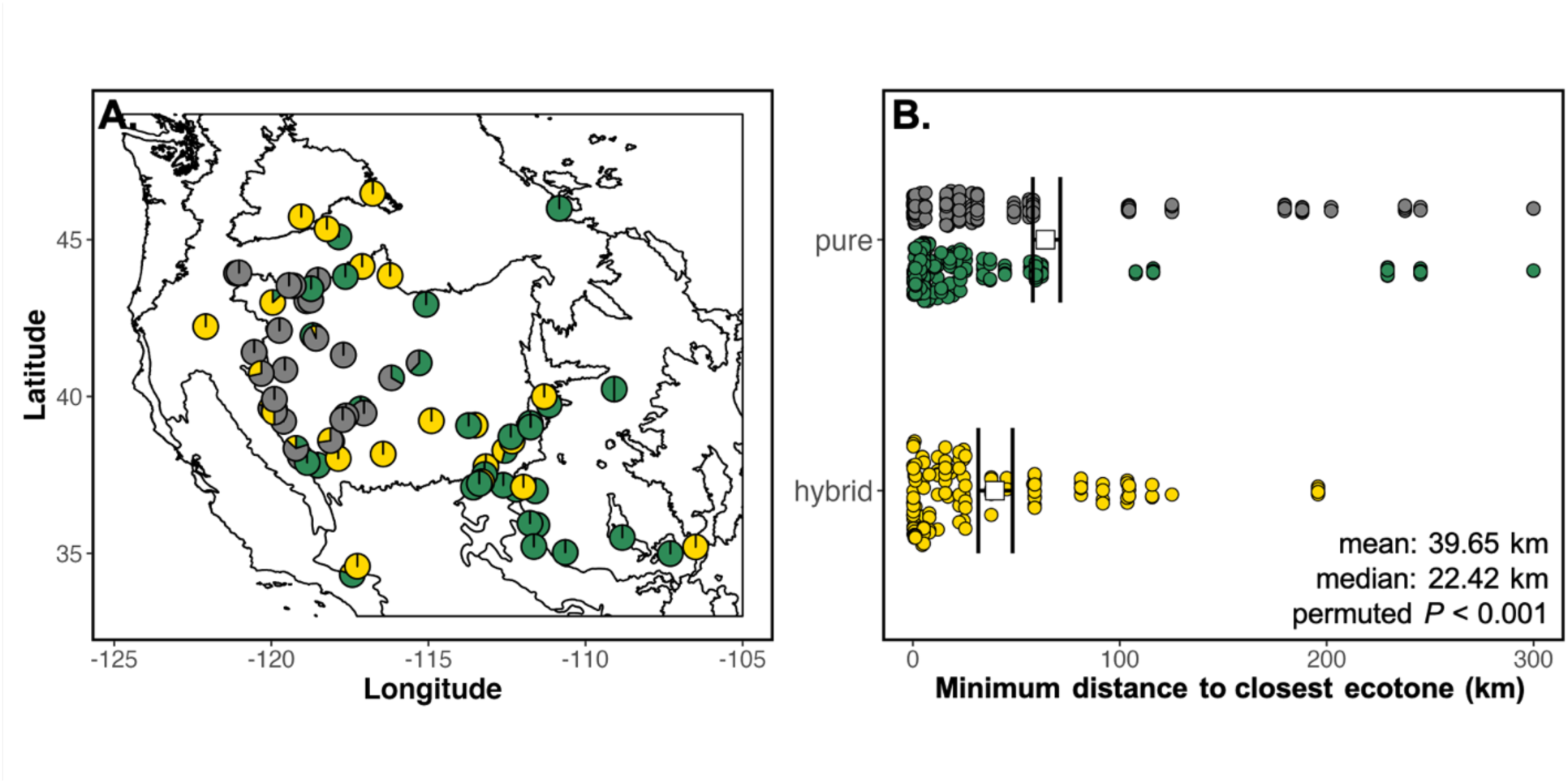
Proximity to ecotone predicts differentiation and hybridization between *E. n. nauseosa* and *E. n. consimilis* lineages. (**A**) Map displaying the proportion of individuals from each population with parental or hybrid ancestry. Solid lines indicate major ecotones between ecoregions (EPA ecoregion level 2) (**B**) Proximity to ecotone predicts likelihood of hybridization. A rarified, permutation test confirmed the estimated mean distance to ecotone for a hybrid was closer, on average, than an individual of pure ancestry. Colored points designate assignment to either *E. n. nauseosa, E. n. consimilis*, or as a hybrid.

Univariate analyses additionally illustrated how environmental and geographic variables were associated with *q* and varied between the lineage categories (Supplementary Figures 4, 5). Longitude was the most strongly variable associated with *q*, the latter of which exhibited a clear west to east pattern for *E. n. nauseosa* and *E. n. consimilis* (*β* = 0.274, *r*^2^ = 0.327). Cumulative AET was also strongly associated with *q* and lineage distinction, increasing dramatically from *E. n. nauseosa* to *E. n. consimilis*, with hybrids being more intermediate (*β* = 0.015, *r*^2^ = 0.162). Slope, AnnTemp, and FallAET were all additionally associated with lineage differentiation, albeit, to a much lesser degree.

Finally, proximity of sampling location to nearest ecotone (designated by ecoregion level 2 boundaries; Omernik & Griffith, 2014) was predictive of hybrid ancestry (hybrid distance: mean = 39.65 km, median = 22.42 km; pure distance: mean = 64.10 km, median = 28.53 km; permuted *P* < 0.001; Figure 4), suggesting a role for ecotone transitions in hybridization. The same environmental association analyses applied to the subsetted dataset with equal sampling across locations and lineages illustrated similar patterns and parameter estimates (Figure S6), confirming that our results were not confounded by uneven sampling effects.

### Assessing the evolutionary cohesiveness of varieties

Phylogenetic reconstruction further illuminated divergence between the two lineages with patterns of substructure coinciding with some varietal designations (Figure 5B; see Supplementary Figure 7 for tree uninfluenced by admixture). High bootstrap support (>75%) characterized many clades within the tree, including: divergence between *E. n. nauseosa* and *E. n. consimilis*, deeper divergences characterizing several southwestern edaphic varieties within *E. n. consimilis*, and monophyly of some populations and varieties. The varieties *bigelovii*, *graveolens*, *juncea*, and *turbinata* were each represented by samples from multiple locations that clustered together monophyletically with respect to each variety (Figure 5B). In contrast, individuals from other varieties appeared in multiple clusters throughout the tree and were represented across both lineages and hybrids (e.g., *oreophila*, *hololeuca*, *salicifolia*, and *speciosa*; Figure 5B). While individuals of hybrid ancestry with respect to *E. n. nauseosa* and *E. n. consimilis* are spread across many varieties, several varieties appear to be of entirely hybrid origin (*mohavensis*, *nana*, and *latisquamea*).

**Figure 5:**
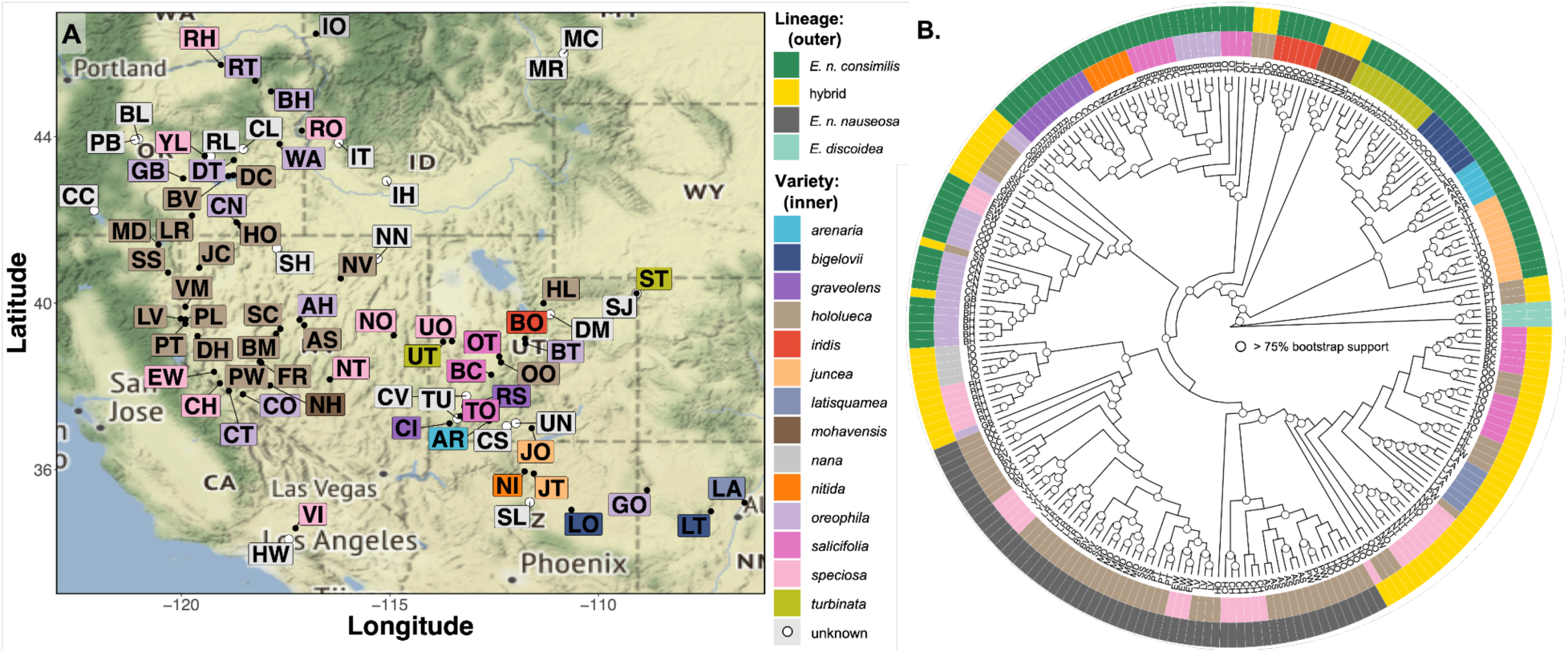
Varietal designation lacks clear evolutionary cohesiveness, likely due to range sympatry. (**A**) Map represents every sampling location within the study colored by the varietal designation. (**B**) Phylogenetic reconstruction of 231 individuals (47 populations, 6 ≤ individuals per population) including *E. discoidea* as the outgroup, built using IQ-TREE. The tip labels are the population IDs and circles at node represent > 75% bootstrap support for the node. The two-colored rings indicate ancestry assignment of lineage or hybrid (outed) and varietal designation (inner).

The digitization of the historic range maps from Anderson (1986b) revealed a large degree of sympatry across the lineages and varieties (Figures 1A, 5A), as well as variation in range size. While the error margins of manually georeferencing hand drawn range maps to digital form should be considered, some inferences of range size and overlap can be useful for this highly complex species. The *E. nauseosa* range was estimated to be ∼2.4mil km^2^, with variety size ranging from just two single locations (*iridis*) to ∼1mil km^2^ (*graveolens*; Figure S8). There is also substantial range overlap of the varieties, specifically concentrated among the edaphic varieties in UT / northern AZ and among varieties in southern CA (Figure S9).

Genetic diversity estimates and inbreeding coefficients (*F*) varied considerably among the lineages and varieties, with the general pattern that the edaphic and narrow endemic varieties in the *E. n. consimilis* lineage had both decreased genetic diversity and increased inbreeding (avg. endemics: *F =* 0.094, *θ_π_*= 0.0185, *θ_W_* = 0.0172; avg. all others: mean *F =* 0.017, *θ_π_*= 0.0196, **θ*_W_* = 0.0202; Figure S10).

## Discussion

Due to complex biogeographic histories and heterogeneous landscapes, numerous groups of plants in the Intermountain West consist of phenotypically and genetically diverged groups with variable reproductive isolation among populations and species with partial sympatry (e.g., Buck et al., 2023; Chhatre et al., 2018; Grant, 1981). Our results indicated that *E. nauseosa* has been strongly affected by the geographic and environmental variation in this region, resulting in two sympatric, independently evolving lineages that largely correspond with the *E. n. nauseosa* and *E. n. consimillis* subspecific designations. Within this distribution, environmental variation predicted the location of subspecific ancestry as well as the occurrence of numerous hybrid populations that occurred in environmentally intermediate locations relative to the two subspecies. Hybrid populations exhibited a continuum of admixed ancestry across a geographically dispersed mosaic hybrid zone, with hybridization more likely in populations closer to ecotone boundaries, which illustrated both replicated instances of hybridization and context dependent variation in hybrid genetic variation. Within each subspecies populations were surprisingly differentiated given the outcrossing mating system and disturbance-oriented nature of this species. While many of the named varieties were phylogenetically dispersed and did not represent evolutionary cohesive groups, a group of southwestern endemic edaphic varieties represented genetically cohesive and differentiated groups on independent evolutionary trajectories. Our study begins to resolve complex patterns of phenotypic and genetic divergence, with ecologically based variation in hybridization, across the distribution of a phenotypically diverse foundational species.

### Genetic differentiation between two subspecific lineages despite sympatric overlap

The changing taxonomy of the *E. nauseosa* complex suggests complex evolutionary history, but historic treatments have all recognized the existence of multiple subspecies, even in sympatry. In support of these observations, the most pronounced feature of genetic variation in *E. nauseosa* we observed was divergence between two lineages that largely coincided with the two recognized subspecies (*E. n. nauseosa* [green] and *E. n. consimillis* [grey]). The geography of divergence is notable - the differentiated lineages occur sympatrically over much of the sampled region and hybrids were found in many geographically distant locations (Figures 1A, 5A, S8, S9; see below) - consistent with past descriptions of the subspecies having largely overlapping distributions (Anderson 1986b, Figure S7). The geography of differentiation documented here suggests the evolution of reproductive isolation between the subspecies. Consistent with this pattern, reports of the subspecies, and even pairs of specific varieties, occurring sympatrically with no sign of trait intergradation or hybridization have been common (Anderson, 1986b; Hanks et al., 1975; Hegerhorst et al., 1987b). In numerous cases of sympatry, the subspecies have been reported to have distinct phytochemical variation and unique communities of gall-forming insects (Floate et al., 1996; Hegerhorst et al., 1987b), suggesting extended ecological consequences of genome divergence among the subspecies.

Population differentiation within each lineage was surprisingly pronounced across spatial and environmental gradients, given insect pollination and wind dispersal of seed. When sample locations with pure ancestry were analyzed separately for each lineage, pronounced patterns of IBD and fine-scale population differentiation were evident (Figure 2, Figure S2). Individuals were almost fully identifiable to sampling location based on genotypic variation alone (Figure 2A, B), as illustrated by Random Forest models which correctly inferred sample location from genotypic data with very little error (individual accuracy = 92.43 – 95.29%). This evidence for differentiation across small distances is consistent with recent analyses of similar data for a smaller set of *E. n. nauseosa* var. *hololeuca* populations (Faske et al., 2021) and suggests that insect pollination and wind-dispersed seed in *E. nauseosa* (which have sepals modified into a pappus, a dispersal mechanism common to many members of the Asteraceae), do not generate substantial gene flow that would otherwise limit local adaptation across environmental gradients (reviewed in Lenormand, 2002). Strong selection and local adaptation to diverse environments could be responsible for maintaining genetic differentiation across small spatial scales, and previous work detected congruent evidence for local adaptation to environmental variation through analyses of genetic environment associations and common garden phenotypic data (Faske et al., 2021). Despite *E. nauseosa* being widespread, strongly outcrossing, and disturbance oriented, genetic differentiation across small spatial scales and local adaptation to diverse environments appear capable of driving fine-scale diversification in this taxonomically diverse species complex.

Many plant groups of western North America have divergence histories associated with Quaternary glacial cycles, including expansions out of refugia and distributional shifts in response to changing climate (Dynesius & Jansson, 2000; Massatti et al., 2018; Massatti & Knowles, 2020; Shafer et al., 2010). Geographic variation of both nucleotide diversity and Tajima’s *D* (Figure 2D, Figure S3) suggest the potential for different histories in the two lineages which could have involved periods of allopatry followed by a progression of increased sympatry. Tajima’s *D* was positively associated with latitude in *E. n. nauseosa* (*r* = 0.274), suggesting a southern expansion, and negatively associated with latitude in *E. n. consimilis* (*r* = -0.494), suggesting a northern expansion (Figure 2D). While negative Tajima’s *D* estimates can arise from selective sweeps, it is unlikely to find this pattern across the whole genome; thus, the patterns here are more likely to reflect demographic changes (Stajich & Hahn, 2005) and could help identify the biogeographic history of these subspecies. One possible scenario is that *E. n. consimilis* expanded from the southern Rocky Mountain region, while *E. n. nauseosa* expanded south from a more northwestern distribution with sympatry occurring after periods of independent evolution. Differences in environmental variation associated with the subspecies (discussed below) may also reflect histories of local adaptation to different conditions. Although outside the scope of work presented here, future analyses spanning a range of demographic models based on more continuous sampling across the distribution would improve our understanding of this history.

### Ecological predictors of hybridization between lineages

While sympatric distributions and the occurrence of individuals with pure *E. n. nauseosa* and *E. n. consimillis* ancestries within the same sampling locations suggest the evolution of reproductive isolation, the occurrence of admixed ancestry in numerous widely dispersed locations illustrates hybridization in a context dependent pattern (i.e., a mosaic hybrid zone; Rand & Harrison, 1989). The general pattern of widely dispersed locations with admixed ancestry across the overlapping distributions of *E. n. nauseosa* and *E. n. consimillis* (Figure 1A, 4A) is consistent with expectations for a mosaic hybrid zone, where reproductive isolation is predicted by ecological and environmental variation rather than a geographic cline of secondary contact (Harrison & Larson, 2016; Rand & Harrison, 1989).

Evidence for environmental variation predicting ancestry (Figure 3A, B) suggests that local adaptation across environmental gradients is likely to mediate reproductive isolation between *E. n. nauseosa* and *E. n. consimilis* or that hybrid fitness varies with environment. The environmental variables most strongly contributing to these results (e.g., CumlAET, FallAET, Slope, PrcpSeas; Figure 3B) are not surprising given similar variables were implicated in recent inference of local adaptation across *E. n. nauseosa* of the western Great Basin (Faske et al., 2021), as well as in other plant taxa of western North America (Massatti & Knowles, 2020; Shryock et al., 2017). Past observations have noted that the timing of flowering and seed-set varies across the range due to environmental conditions (Meyer et al., 1989), and it has been anecdotally observed that these traits can differ between *E. n. nauseosa* and *E. n. consimillis* when in sympatry, suggesting that local adaptation may also underlie prezygotic isolation.

Admixed ancestries were also more likely to occur near ecological transitions than were non-hybrid ancestries (Figure 4). This is not surprising, as environment and climate transition across the North American ecoregion level 2 boundaries on which we based analyses (Omernik & Griffith, 2014), but suggests that ecotones could additionally predict the distribution of hybrids. Many of the ecoregion 2 boundaries cover biogeographic breaks that have been described as suture zones, where numerous taxa hybridize at overlapping secondary contact zones (Remington, 1968; Swenson & Howard, 2004, 2005). Our results suggest that the multiple suture zones spanned by the large, biogeographically complex distribution of *E. nauseosa* contribute to the folding of ancestry across intermediate environments.

The decoupling of environment and geography is a feature of our analyses (Figure S2) that is shared with other studies of mosaic hybrid zones (e.g., *Bombina*, Vines et al., 2003; *Populus*, Lindtke et al., 2013; *Gryllus*, Ross & Harrison, 2002; *Mytilus*, Bierne et al., 2003). Such systems are valuable for identifying the mechanisms that underlie the repeated and independent formation of hybrids, but may often illustrate context dependence and stochasticity in admixture among populations and across genomes (Mandeville et al., 2017; Teeter et al., 2010). Given our ability to predict the location of admixed populations based on both environmental variation (Figure 3D) and proximity to ecological transitions (Figure 4), one might expect a similar genomic composition for hybrids if the environment serves as a strong filter for hybrids relative to non-admixed individuals. We found a contrasting pattern in which hybrids were found throughout RDA space, indicating substantial variation in hybrid genetic variation and its environmental associations (Figure 3B). This stochasticity suggests the interaction between environmental variation and hybrid fitness varies across the range of *E. nauseosa*, which has also been reported in other species inhabiting diverse environments (e.g., Lexer et al., 2004; Mandeville et al., 2017; McFarlane et al., 2022). As hybrids can have phenotypic variation distinct from either parent species (Kagawa & Takimoto, 2018; Rieseberg et al., 1999, 2007), this could also suggest that hybridization may generate genomic variation that can effectively explore the environmental fitness landscapes occupied and unoccupied by the parental lineages.

### The evolutionary cohesiveness of varieties

Reconstruction of the evolutionary relationships among samples spanning different named varieties revealed a spectrum of outcomes, ranging from monophyly, consistent with a variety being an evolutionarily cohesive lineage, to polyphyly, where individuals from the same putative variety were found throughout the phylogeny. Several varieties, including *speciosa*, *salicifolia*, *oreophila,* and *hololeuca*, showed instances of polyphyly (Figures 1A, 5A). Varietal designation in such cases could be a result of phenotypic similarity of distantly related populations, which could arise through phenotypic plasticity and/or convergent evolution to similar ecological conditions. Both processes can challenge taxonomy and have been widely documented in plant taxa of western North America (e.g., Love et al., 2023; Wilson et al., 2007). Indeed, difficulties in identifying field collected *E. nauseosa* to variety could obfuscate our analyses and challenge our ability to evaluate the evolutionary cohesiveness of some varieties. For example, Anderson (1986b) suggested that the *hololeucus* and *oreophila* varieties maintained phenotypic differentiation in sympatry to the extent they could be recognized as different species, which contrasts with polyphyly of these varieties observed here. However, *hololeuca* and *speciosa* both had polyphyletic populations with admixed subspecific ancestry, suggesting the mosaic distribution of hybridization described above could also have influenced these analyses. Nonetheless, the numerous occurrences of polyphyly reported here suggest much future work will be needed to resolve the varietal level taxonomy of rubber rabbitbrush.

In contrast, some groups illustrated genetic variation consistent with varietal designations. For instance, vars. *bigelovii*, *graveolens*, and *turbinata* were all cohesive, monophyletic groupings of samples spanning multiple locations. The other seemingly monophyletic variety from multiple locations is *juncea*, which along with *arenaria*, *bigelovii*, and *turbinata* formed a distinct, substructured group within the *E. n. consimillis* lineage. These varieties, many of which had reduced genetic diversity (Fig. S10), are largely thought to be edaphic endemics (Cronquist, 1994) and are primarily found within the southeastern most part of the range (UT and northern AZ). This region coincidentally sees the most amount of varietal overlap (Figures 5A, S8, S9) and contains a large portion of the sampling locations with admixed ancestry with respect to subspecies. Several endemic varieties that formed monophyletic groups (*mohavensis*, *nana*, and *latisquamea*) consisted entirely of admixed ancestry from the two subspecies (Figure 1, Figure 5). As hybrids often have distinct, or transgressive, phenotypic variation with respect to parent species (Kagawa & Takimoto, 2018; Rieseberg et al., 1999, 2007), this result suggests that hybridization could facilitate the colonization of, and local adaptation to, novel environments in these narrow endemic and phenotypically distinct varieties. While this result hints at a role for admixture in varietal diversification, our sampling of individuals and locations is limited for these varieties and future work with more comprehensive sampling will support a more thorough understanding of the potential for hybridization to provide a creative role in the evolution of morphological novelty.

## Conclusions

Our analyses illustrate a continuum of divergence across lineages and environments occupied by *E. nauseosa*, a foundational shrub species of the intermountain west known for its marked phenotypic variation. Our analyses indicate that *E. nauseosa* comprises two genetically distinct lineages that exhibit environmentally dependent reproductive isolation and maintain differentiation despite widely overlapping distributions. We found admixed ancestries in a subset of locations that were widely distributed in a mosaic hybrid zone where ancestry was strongly predicted by both climate variation and the proximity to ecotonal transitions. While most studies of hybridization focus on one instance of secondary contact, those that have looked at multiple zones of contact have found mixed evidence of predictability in the occurrence and outcome of hybridization. Our results suggest that environmental variation is a key factor in mediating reproductive isolation in these young lineages, but also illustrate considerable variation in the association of genetic and environmental variation across multiple instances of hybridization. This is the first geographically extensive survey of genetic variation in *E. nauseosa*, yet we lacked sampling in the southeastern and northeastern portions of the distribution, and our sampling of putative named varieties was limited in scope and space. Inference of evolutionary history for many of the named varieties we sampled may be confounded by potential for misidentification of field collected samples as well as our sampling of only one or two localities across small geographic distances in a species with substantial population structure. Future analyses with more comprehensive sampling of the range, including sampling of many locations within specific named varieties, stand to further shape our understanding of how environment, history, and admixture underlie the extensive phenotypic variation the group is noted for.

## Author Contributions

EAL, BAR, and TLP developed the original idea for this research and secured funding. ACA and BAR (with associated field crews) collected samples and identified them to subspecies / variety. LMS extracted the DNA and prepped the libraries for Illumina sequencing. TMF analyzed the dataset with COM conducting the phylogenetic analyses. TMF, JPJ, and TLP wrote the initial draft and all authors contributed to revising the manuscript.

## Data Availability

The data that supports the finding of this study are available from the Dryad Digital Repository: doi:10.5061/dryad.5hqbzkhbg. This includes population-level summaries of environmental data, information of sampling locality, filtered VCF file, and a matrix of genotype probabilities. All scripts from pre-processing, variant calling, to statistical analyses and data visualization are available on either Dyrad or GitHub: https://github.com/trevorfaske/ERNA_analyses.

