## Supplemental text for "Environment predicts the maintenance of reproductive isolation in a mosaic hybrid zone of rubber rabbitbrush"

### Methods

*Genetic diversity and individual inbreeding coefficient estimation*

We estimated nucleotide diversity ($\theta_\pi$; Nei & Li, 1979)and the Watterson estimator ($\theta_W$; Watterson, 1975) within each population (min. 3 individuals) using methods that incorporate genotype uncertainty implemented in ANGSD V 0.923 (Korneliussen et al., 2013, 2014). Using individual bam files, we estimated the folded site allele frequency likelihoods with "realsfs" and calculated genotype likelihoods using the setting "GL 1" estimated from the model implemented in SAMTOOLS model to obtain the likelihood of the folded site frequency spectrum (SFS). We estimated diversity ($\theta_\pi$ and $\theta_W$) and Tajima's *D* (Tajima, 1989) using "doThetas 1" and "thetastat" commands using SFS likelihoods as priors, respectively, for each loci across the reference assembly and averaged the measures for each population. Additionally, individual inbreeding coefficients (*F*) were calculated while incorporating genotype uncertainty using NGSF (Vieira et al., 2016), implemented through ANGSD-WRAPPER V 0.933 (Durvasula et al., 2016). First, genotype likelihoods were estimated using ANGSD and mapped to the reference assembly with a minimum bp quality of 20 and mapping quality of 30. In order to speed up convergence and account for the inherent low coverage of the data, we use the approximate expectation-maximum (EM) algorithm within NGSF and a minimum ε of $1e^{-8}$.

*Genetic-environment-landscape associations*

**Aim 1:** Redundancy analysis (RDA) was used to show how genetic variation between individuals (i.e., genotype probabilities) was driven by the 10 environmental variables and the inclusion of geographic variation (i.e., latitude/longitude; Figure 3B). Additionally, a variance partitioning technique (Borcard et al., 1992) was used to estimate individual and shared contributions of geography and environment of ancestry coefficients (*q*). This approach uses partial RDAs to estimate the proportion of explained variance for each predictor variable, independently and combined, out of the total explained variance (Figure 3C). All analyses and data manipulation were performed using the base STATS library or custom R functions (R Core Team, 2020), except the library VEGAN (Oksanen et al., 2019) and the *varpart*() function were used for the RDA and variance partitioning. **Aim 2:** To assess the predictability of lineage and hybrid assignment from environmental data, Random Forest models (Breiman, 2001) were built with individual ancestry coefficients (*q*) as the response and environmental data as predictors. Models were trained with 75% of the data and predicted the other 25% test data across $n = 1000$ permutations. Model performance was assessed using $r^2$, root mean square error (RMSE), and a confusion matrix based on the actual versus predicted lineage classification (*E. n. nauseosa* $< 0.08 \leq$ hybrid $\leq 0.87 <$ *E. n. consimilis*) (Figure 3D). **Aim 3:** Univariate models were used to characterize environmental and geographic predictors of ancestry (*q)* and lineage assignment for each population / lineage combination (Figure S6, S7). Beta regressions (BETAREG package in R; Cribari-Neto & Zeileis, 2010) were used for models where *q* was the response because it is bound between 0 – 1, whereas type III ANOVAs (CAR package in R; Fox & Weisberg, 2019) were used for models where lineage assignment was the response. All univariate models were weighted by the number of individuals.

*Hybridization predicted by distance to ecotone*

Following classic hybrid zone theory, the probability of hybridization among our sampled populations occurred at a higher probability with increased proximity to ecotones (i.e., changes from one ecoregion to the next). Distance from individual locality to the nearest ecotone was estimated using EPA guided North American ecoregion levels 2 (Omernik & Griffith, 2014). To control for potential influences of sampling, a permutation-based test (*n* = 10000) was implemented, randomly assigning individuals as either lineage or hybrid at equal numbers in the data and estimating the mean minimum geographic distance to an ecoregion boundary (Figure 4). All analyses were conducted separately for both ecoregion levels and performed using the LME4 library (Bates et al., 2015) or custom R functions.

### Supplemental figures

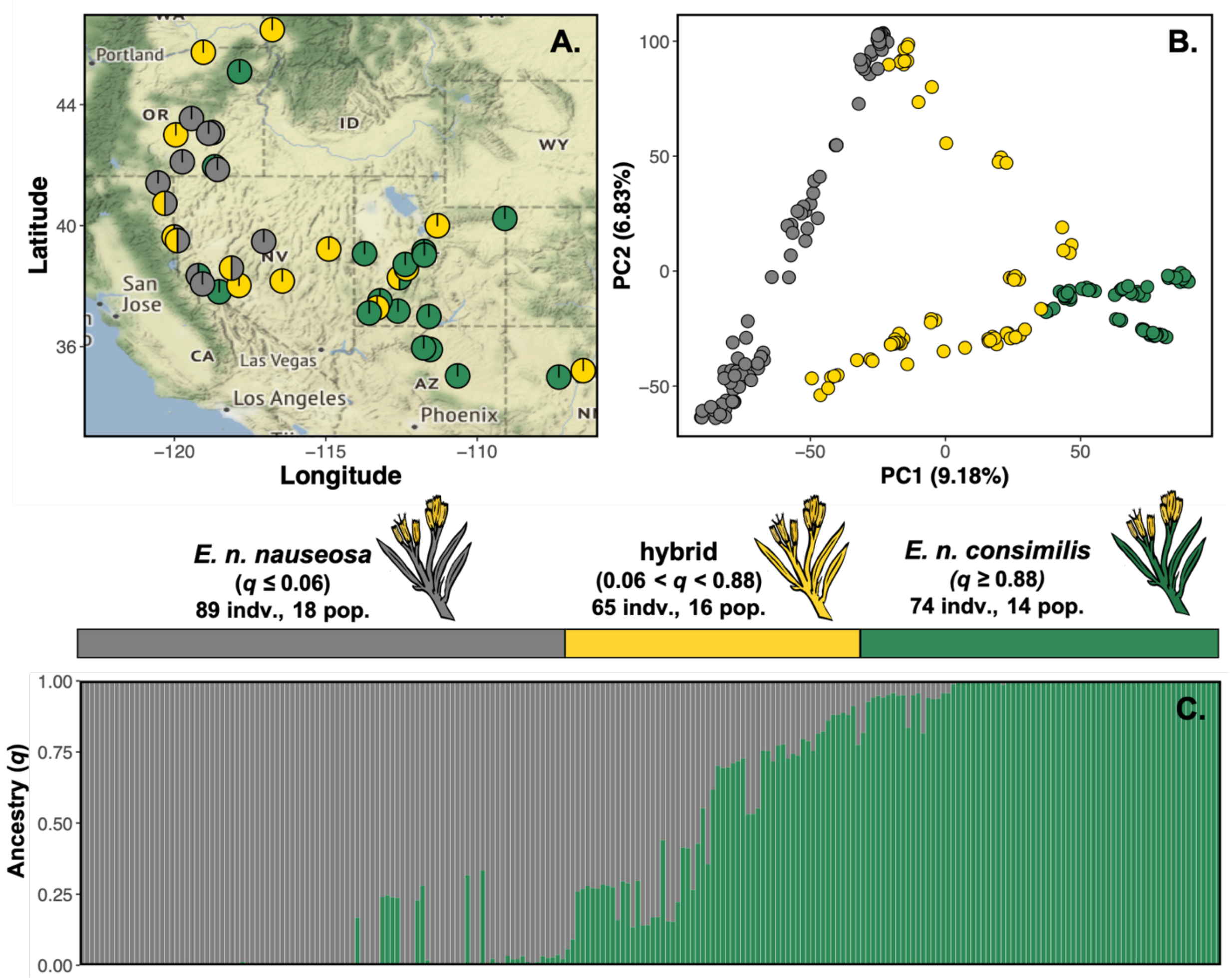


**Figure S1: Analyses of the subsetted and rarified dataset depict the same patterns of genetic differentiation and admixture among lineages and as identical analysis of the full dataset.** (**A**) Sympatric ranges of *E. n. nauseosa* and *E. n. consimilis* based on taxonomic classification in Anderson (1986b) overlaid with sampling locations. Pie chart colors correspond to the proportion of individuals at a locality assigned to each lineage or as a hybrid. (**B**) Principal component analysis (PCA) illustrates genetic differentiation between lineages with the presence of hybrid individuals. Circles represent mean PC values for populations, pie charts reflect ancestry, and lines connect PC values for individuals to population means. Yellow lines represent hybrid individuals with ancestry coefficients $0.06 < q < 0.87$. (**C**) Ancestry coefficients ($q$) from the hierarchical Bayesian model of entropy of $k = 2$ for each individual, ordered by PC1. Color proportion of each bar ancestry proportion.

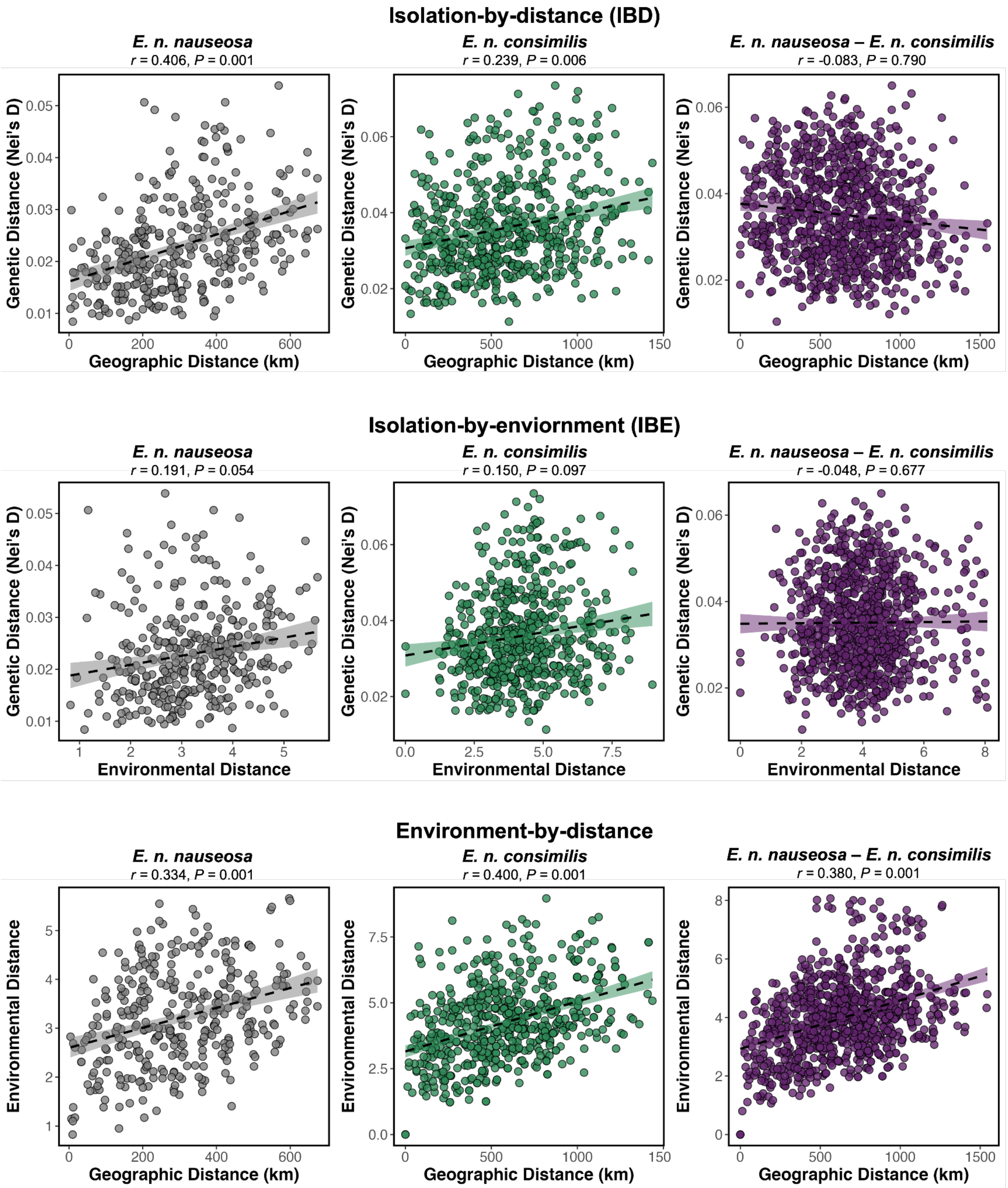


**Figure S2: Strong patterns of isolation-by-distance (IBD) and isolation-by-environment (IBE) within *E. n. nauseosa* and *E. n. consimilis* but not when comparing populations across lineages (shown in purple) although population differentiation remains high.** Raw data format of figures 2C and 3A from the main text, with the addition of the relationship between geographic and environmental distance.  also largely corresponded to geographic distance across the sampled range.  The fitted lines and standard error were estimated using the stat_smooth() function in GGPLOT2 in R. Correlations (*r*) and *P*-values are reported above each panel and were assessed using Mantel tests.

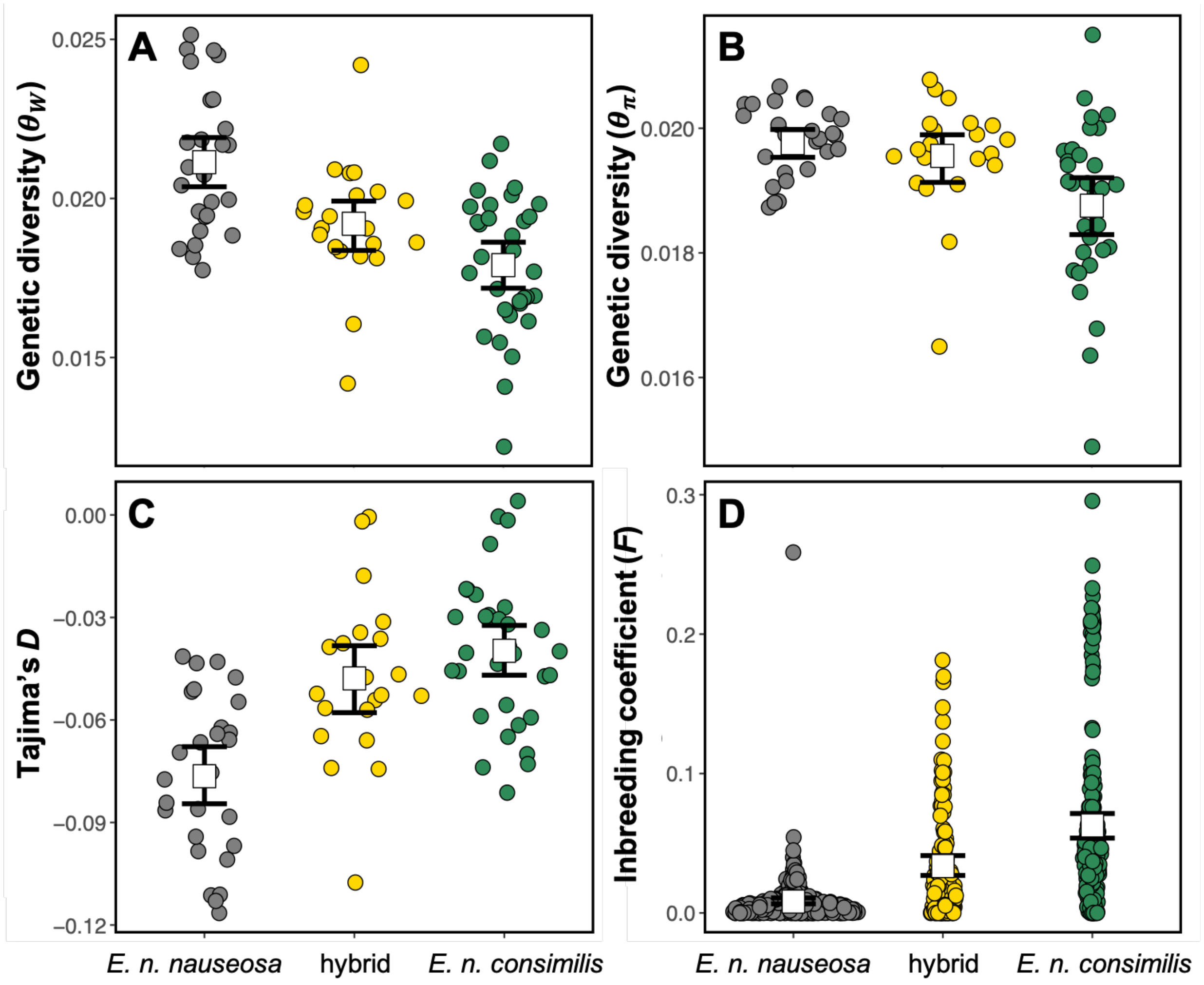


**Figure S3: Genetic diversity, Tajima's *D*, and inbreeding coefficients vary between lineages and hybrids.** (**A**) and (**B**) Genetic diversity (measured by $\theta_W$ and $\theta_\pi$) is greatest in *E. n. nauseosa* and least in *E. n. consimilis,* with hybrids exhibiting intermediate diversity, as expected. (**C**) Lineages also vary in genome-wide Tajima's *D*, with *E. n. consimilis* being the greatest, *E. n. nauseosa* being least, and again, hybrids in-between. (**D**) Similarly, individual inbreeding coefficients (*F*) vary across the lineages and the hybrids, but are generally low. The increase in inbreeding within *E. n. consimilis* is likely due to this lineage containing the edaphic endemics (see Figure S8). The circles represent the raw data while white squares represent the mean and error bars are the 95% bootstrapped confidence interval.

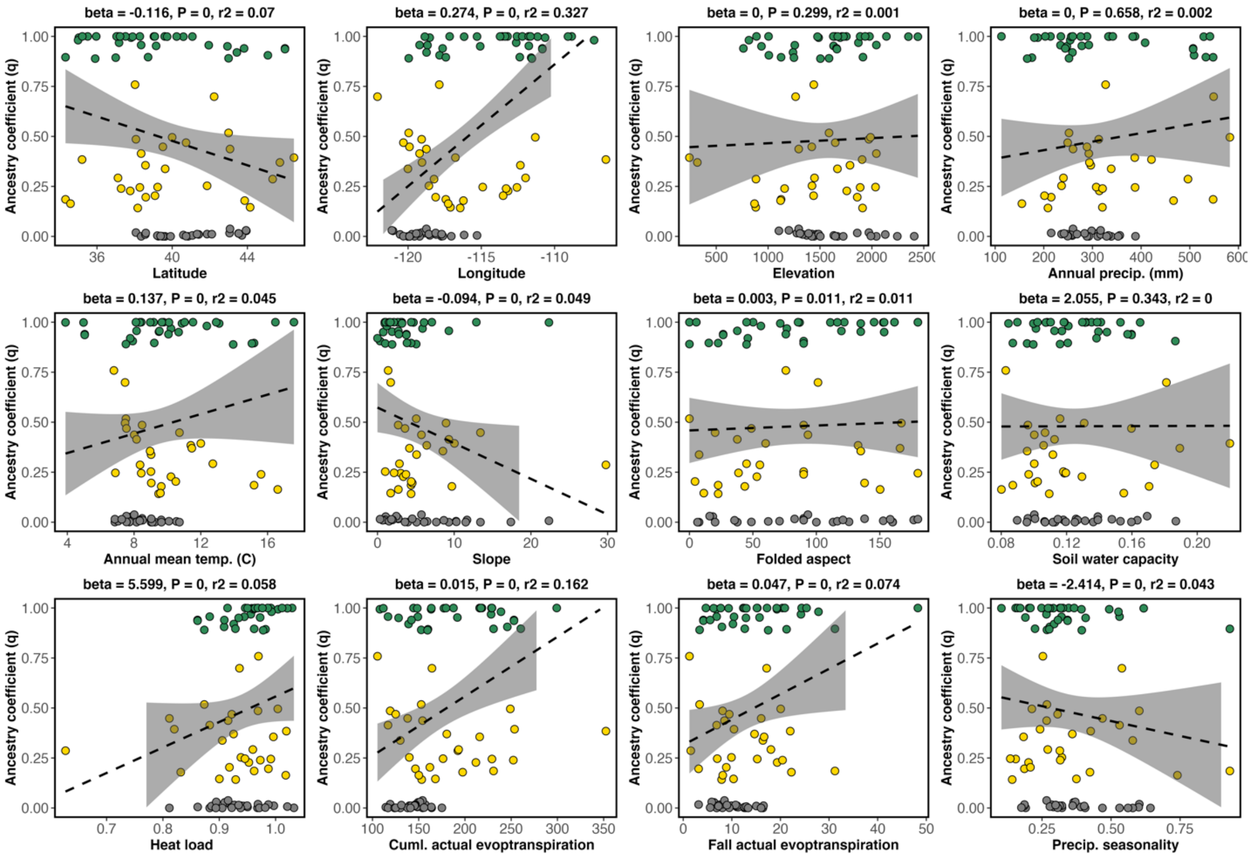


**Figure S4: Association of ancestry coefficients (*q*) with each geographic and environmental variable.** Univariate scatter plots of each variable to ancestry coefficients with fitted line and standard error estimated using the stat_smooth() function in GGPLOT2 in R. Colors correspond to ancestral classification of each individual. As ancestry coefficients are bound from 0 to 1, beta regressions were used to assess the relationship between *q* and predictor variables with beta coefficients, *P*-values, and $r^2$ reported above each panel.

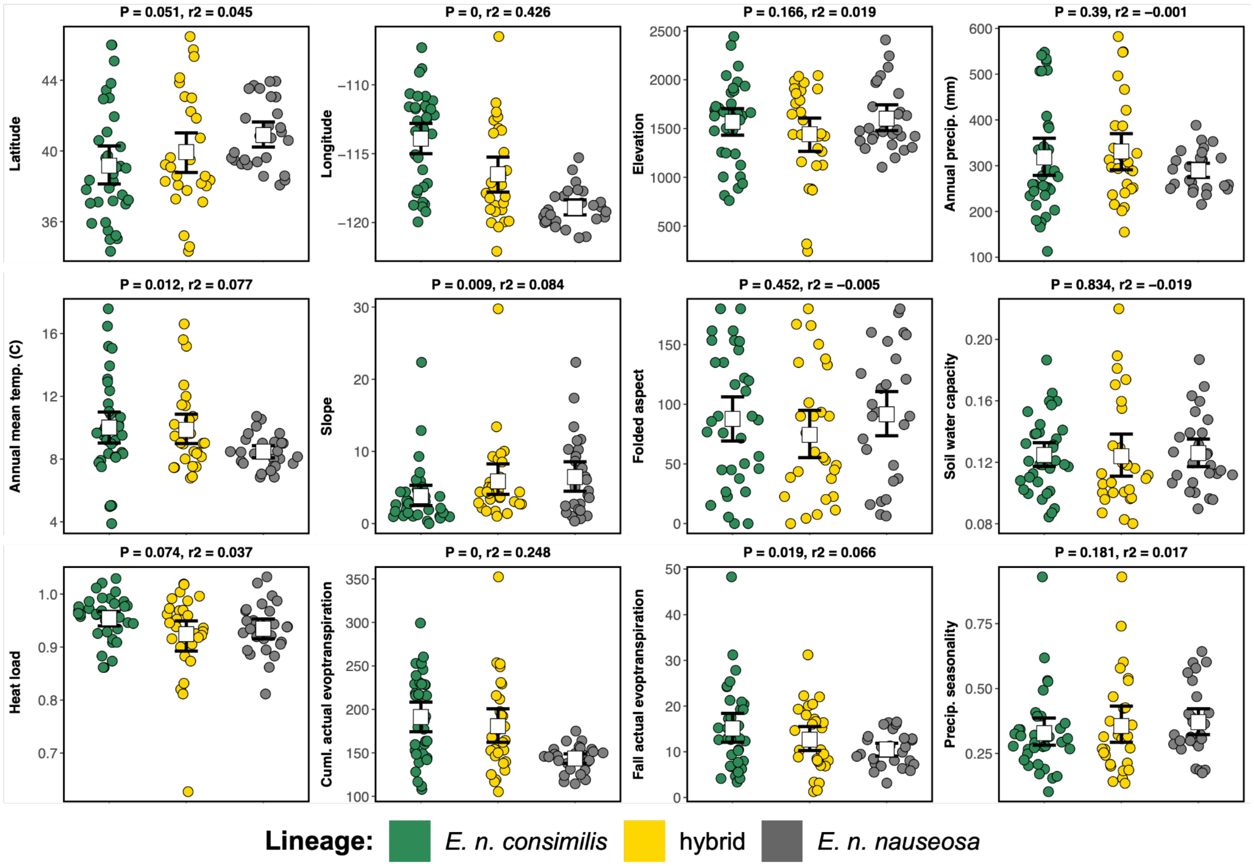


**Figure S5: Geographic and environmental variation varies by ancestral categorization.** Each panel represents a different geographic or environmental variable grouped by either lineage or hybrid. Circles represent the raw data colored by lineage or hybrids while white squares and error bars represent the mean and the 95% bootstrapped confidence interval. *P*-values and $r^2$ were assessed using univariate ANOVA (type III) and are reported above each panel.

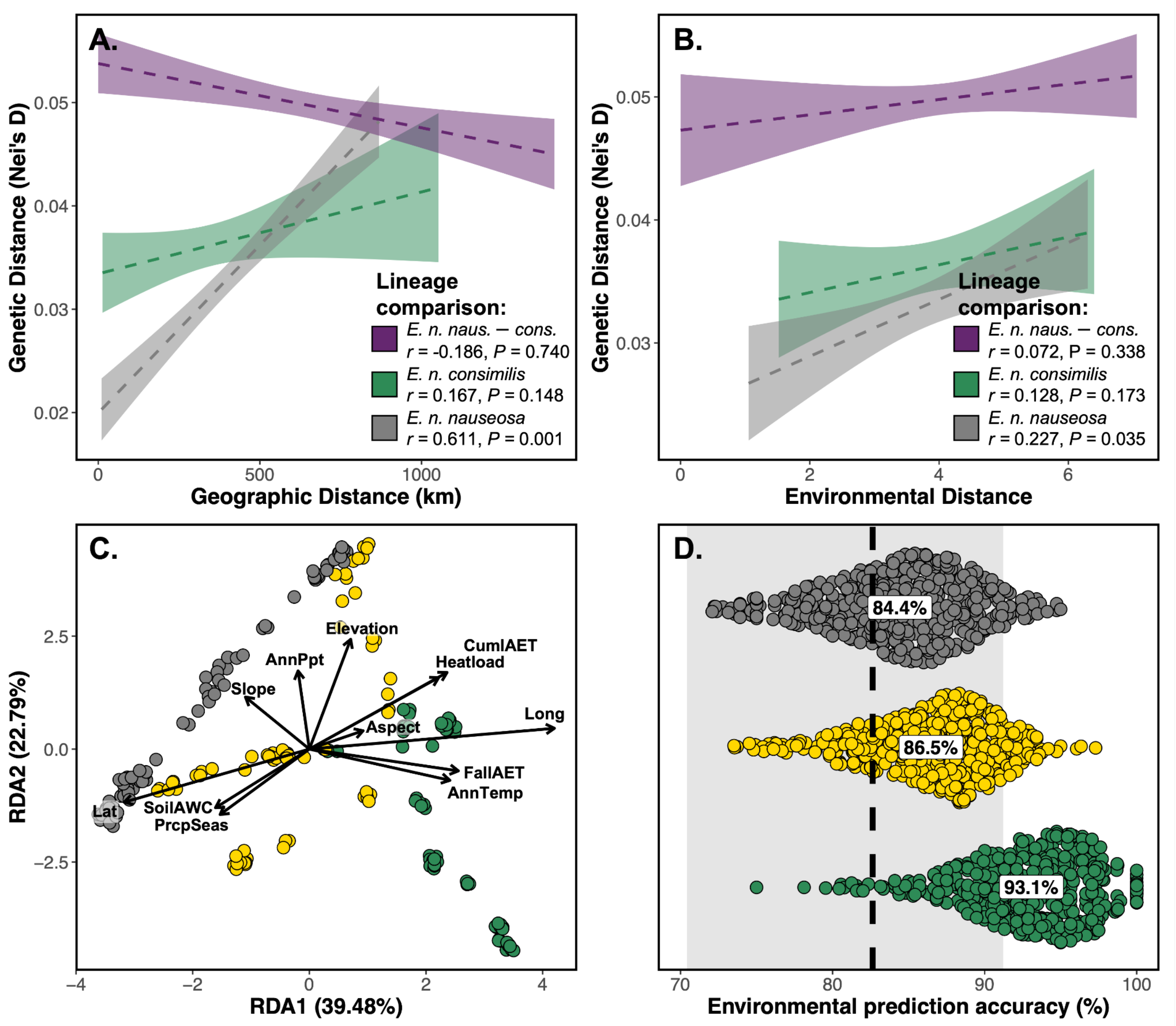


**Figure S6: Analyses of the subsetted and rarified dataset detected the same patterns of geographic and environmental variables on genetic variation as identical analyses of the full full dataset.** (**A**) Strong patterns of isolation-by-distance (IBD) were present within the two lineages but not when comparing populations between lineages (shown in purple). Figure 2C in main text. (**B**) Strong patterns of isolation-by-environment (IBE) were present within the two lineages but not when comparing populations between lineages (shown in purple). Figure 3A in main text. (**C**) Environmental variation associated with genetic structure between the lineages illustrated by RDA. Axes were rotated in order to match that of the PCA in Figure 1 and environmental loadings were scaled (4.36×) for graphical representation. Figure 3B in main text. (**D**) Environment alone predicts an individuals' classification to either lineage or as a hybrid using Random Forest. Points represent permuted accuracy within each lineage with means for each shown in text. Dashed line and shaded bar represent the overall mean predictive accuracy (82.6%) and 95% confidence interval. Figure 3D in main text.

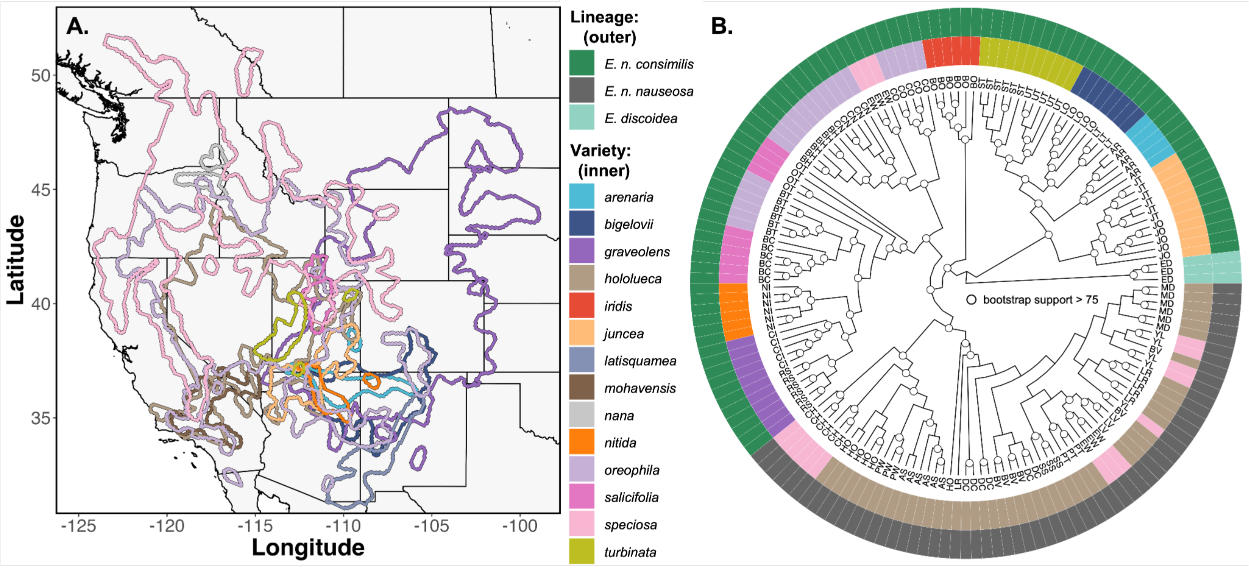


**Figure S7: Phylogeny with no hybrids shows monophyletic split of *E. n. nauseosa* and *E. n. consimilis*.** (**A**) Lines representing the boundaries of digitized range maps from Anderson (1986b) for the varieties sampled in our study. Extensive overlap highlights the degree of sympatry and variation in range size illustrates complexity of varietal taxonomic designation. (**B**) Phylogenetic reconstruction of same tree depicted in Figure 5B of the main test with hybrids removed, built using IQ-TREE. The tip labels are the population IDs and circles at node represent > 75% bootstrap support for the node. The two-colored rings indicate ancestry assignment of lineage or hybrid (outed) and varietal designation (inner).

| | graveolens | oreophila | speciosa | hololeuca | bigelovii | juncea | latisquamea | mohavensis | arenaria | turbinata | salicifolia | nitida | nana | iridis |
|---|---|---|---|---|---|---|---|---|---|---|---|---|---|---|
| graveolens 1018913 | 100 | 16.5 | 8.2 | 7.3 | 15 | 6.1 | 0.9 | 0 | 5.8 | 1 | 1.8 | 2.9 | 0 | 0 |
| oreophila 882012 | 19.1 | 100 | 37.4 | 49.2 | 5.8 | 2.6 | 0.1 | 3.9 | 0.6 | 4.4 | 3 | 0.8 | 0.3 | 0 |
| speciosa 752439 | 11.1 | 43.8 | 100 | 14.2 | 0 | 0 | 0 | 1.6 | 0 | 0.1 | 3.4 | 0 | 3 | 0 |
| hololeuca 504797 | 14.6 | 86 | 21.2 | 100 | 1.2 | 4.5 | 0 | 9 | 1.1 | 7.5 | 4 | 0.8 | 0 | 0 |
| bigelovii 175978 | 86.6 | 29.3 | 0 | 3.5 | 100 | 7.6 | 6.8 | 0 | 25.1 | 0 | 0 | 14.4 | 0 | 0 |
| juncea 81562 | 76 | 28.4 | 0 | 27.9 | 16.5 | 100 | 0 | 0 | 14.1 | 1.5 | 0 | 12 | 0 | 0 |
| latisquamea 74685 | 12.7 | 1.1 | 0 | 0 | 16 | 0 | 100 | 0 | 0 | 0 | 0 | 0 | 0 | 0 |
| mohavensis 65944 | 0 | 52.7 | 18.3 | 68.7 | 0 | 0 | 0 | 100 | 0 | 0 | 0 | 0 | 0 | 0 |
| arenaria 58617 | 100 | 8.4 | 0 | 9.5 | 75.3 | 19.6 | 0 | 0 | 100 | 3.9 | 0 | 27.7 | 0 | 0 |
| turbinata 39785 | 26.8 | 97.6 | 1.5 | 94.7 | 0 | 3.1 | 0 | 0 | 5.8 | 100 | 0 | 2.3 | 0 | 0 |
| salicifolia 29887 | 60.2 | 89 | 86 | 66.9 | 0 | 0 | 0 | 0 | 0 | 0 | 100 | 0 | 0 | 0 |
| nitida 29822 | 99.8 | 23.9 | 0 | 12.9 | 84.9 | 32.8 | 0 | 0 | 54.5 | 3 | 0 | 100 | 0 | 0 |
| nana 22303 | 0 | 12.4 | 100 | 0 | 0 | 0 | 0 | 0 | 0 | 0 | 0 | 0 | 100 | 0 |
| iridis 2 | 100 | 100 | 0 | 100 | 0 | 0 | 0 | 0 | 0 | 0 | 100 | 0 | 0 | 100 |

**Figure S8: Substantial range overlap and variation in range size among the sampled varietal designations.** Pairwise percent range overlap of each variety across columns. Area in km$^2$ under each variety. Presence of color represents any amount of overlap while gradient of color is proportional to amount out of the total area of that variety. Varieties are ordered by decreasing range size.

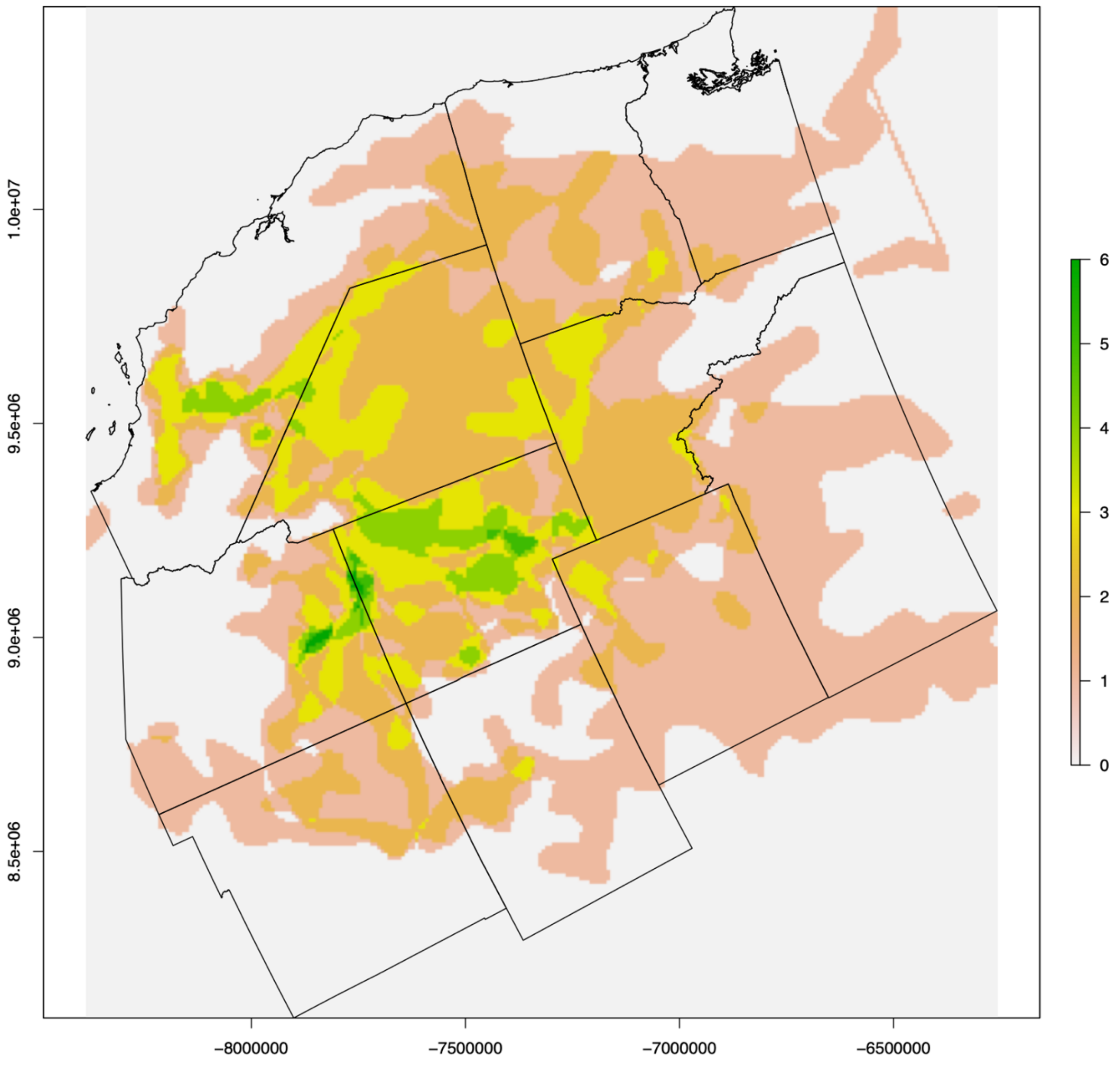


**Figure S9: Substantial range overlap among the sampled varietal designations.** Map highlights the range of all varieties and the color ramp corresponds to the number of overlapping varieties in a given region (max. = 6).

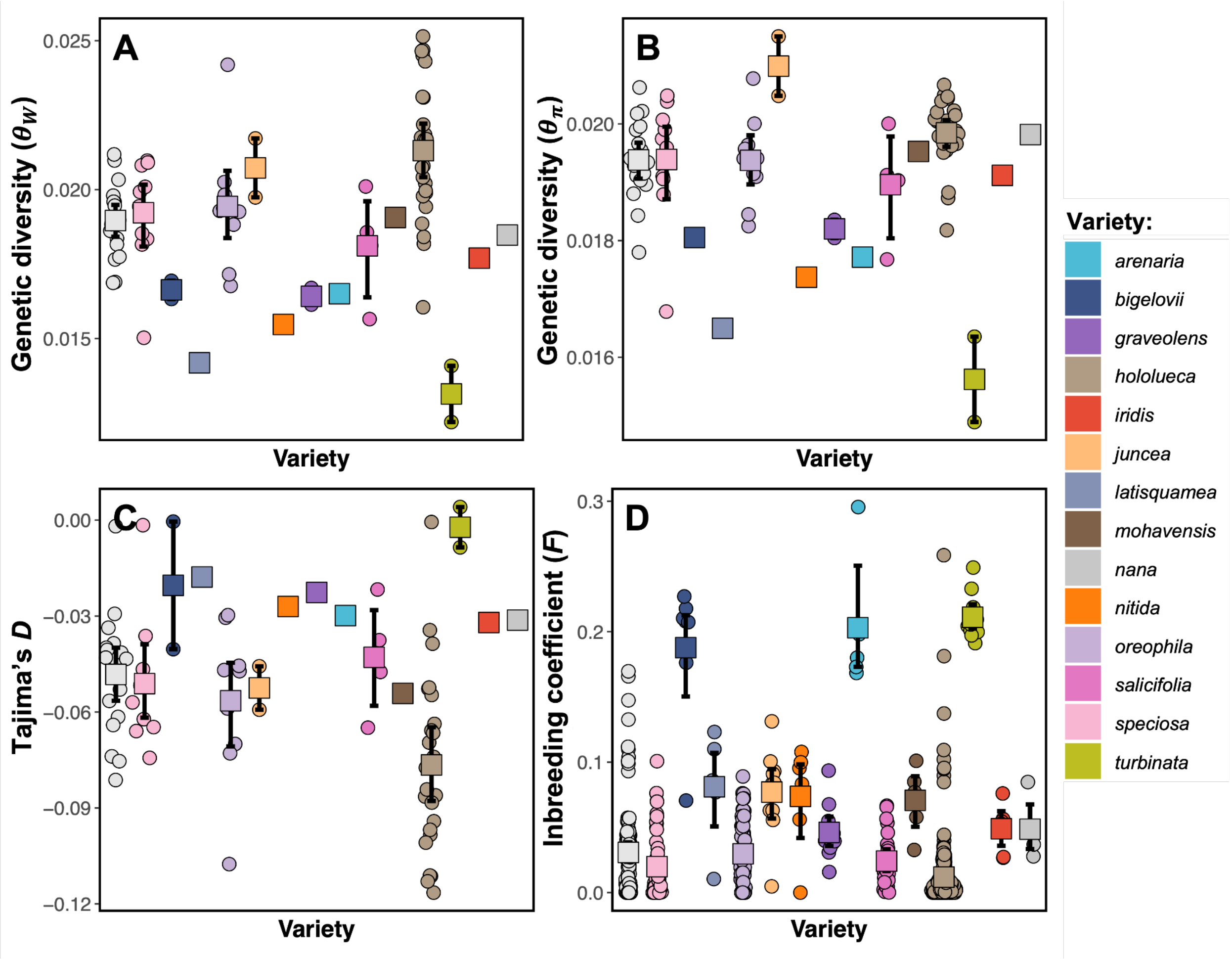


**Figure S10: Genetic diversity, Tajima's *D*, and inbreeding coefficients vary across varietal designations with more edaphic, endemic varieties having decreased diversity and increased inbreeding.** (**A**) and (**B**) Genetic diversity (measured by $\theta_W$ and $\theta_\pi$), (**C**) genome-wide Tajima's *D*, and (**D**) individual inbreeding coefficients (*F*) across varieties. Varieties are ordered by mean latitude of sampling locations. The circles represent the raw data while white squares represent the mean and error bars are the 95% bootstrapped confidence interval.

**Table S1: Information on sampling location and ancestry classification for each.** Pop: 2 letter sampling abbreviation; Elev: elevation in meters; Anc: mean ancestry coefficient estimated from ENTROPY $k = 2$ model; *consimilis,* hybrid, *nauseosa*: number of individuals assigned to each of the lineage designations; N: sample size for each location.

| Pop | Latitude | Longitude | Elev. | Variety | Anc | *consimilis* | hybrid | *nauseosa* | N |
|---|---|---|---|---|---|---|---|---|---|
| AH | 39.601 | -117.160 | 1754 | *oreophila* | 1.000 | 15 | 0 | 0 | 15 |
| AR | 37.174 | -112.618 | 1870 | *arenaria* | 0.968 | 5 | 0 | 0 | 5 |
| AS | 39.473 | -117.049 | 2408 | *hololeuca* | 0.001 | 0 | 0 | 15 | 15 |
| BC | 38.280 | -112.568 | 1887 | *salicifolia* | 0.651 | 7 | 6 | 0 | 13 |
| BH | 45.089 | -117.848 | 1007 | *oreophila* | 0.906 | 6 | 0 | 0 | 6 |
| BL | 43.944 | -121.028 | 1104 | unknown | 0.028 | 0 | 0 | 6 | 6 |
| BM | 39.264 | -117.724 | 2245 | *hololeuca* | 0.001 | 0 | 0 | 14 | 14 |
| BO | 39.145 | -111.756 | 1631 | *iridis* | 0.952 | 6 | 0 | 0 | 6 |
| BT | 39.028 | -111.746 | 1711 | *oreophila* | 1.000 | 6 | 0 | 0 | 6 |
| BV | 43.053 | -118.872 | 1277 | *hololeuca* | 0.016 | 0 | 0 | 15 | 15 |
| CC | 42.231 | -122.087 | 1265 | unknown | 0.699 | 0 | 3 | 0 | 3 |
| CH | 38.072 | -119.073 | 1971 | *speciosa* | 0.086 | 0 | 1 | 6 | 7 |
| CI | 37.112 | -113.567 | 888 | *graveolens* | 1.000 | 5 | 0 | 0 | 5 |
| CL | 43.708 | -118.516 | 1372 | unknown | 0.005 | 0 | 0 | 6 | 6 |
| CN | 41.945 | -118.672 | 1257 | *oreophila* | 0.956 | 5 | 0 | 0 | 5 |
| CO | 37.811 | -118.521 | 1694 | *oreophila* | 0.994 | 5 | 0 | 0 | 5 |
| CS | 37.043 | -112.199 | 1647 | unknown | 1.000 | 6 | 0 | 0 | 6 |
| CT | 37.895 | -118.857 | 2443 | *oreophila* | 0.999 | 6 | 0 | 0 | 6 |
| CV | 37.767 | -113.169 | 1758 | unknown | 0.228 | 0 | 6 | 0 | 6 |
| DC | 43.076 | -118.749 | 1295 | *hololeuca* | 0.064 | 0 | 1 | 14 | 15 |
| DH | 39.211 | -119.608 | 1423 | *hololeuca* | 0.004 | 0 | 0 | 15 | 15 |
| DM | 39.731 | -111.156 | 2352 | unknown | 0.999 | 4 | 0 | 0 | 4 |
| DT | 43.439 | -118.738 | 1240 | *oreophila* | 0.920 | 6 | 0 | 0 | 6 |
| EW | 38.348 | -119.212 | 2043 | *speciosa* | 0.247 | 3 | 2 | 10 | 15 |
| FR | 38.571 | -118.082 | 2129 | *hololeuca* | 0.010 | 0 | 0 | 15 | 15 |
| GB | 42.999 | -119.954 | 1587 | *oreophila* | 0.567 | 2 | 13 | 0 | 15 |
| GO | 35.505 | -108.829 | 1977 | *oreophila* | 1.000 | 6 | 0 | 0 | 6 |
| HL | 39.997 | -111.314 | 1987 | *hololeuca* | 0.496 | 0 | 5 | 0 | 5 |
| HO | 41.855 | -118.577 | 1423 | *hololeuca* | 0.028 | 0 | 1 | 14 | 15 |
| HW | 34.325 | -117.422 | 1125 | unknown | 0.754 | 4 | 1 | 0 | 5 |
| IH | 42.938 | -115.075 | 936 | unknown | 0.950 | 5 | 0 | 0 | 5 |
| IO | 46.475 | -116.769 | 242 | *nana* | 0.394 | 0 | 5 | 0 | 5 |
| IT | 43.856 | -116.220 | 1120 | unknown | 0.179 | 0 | 5 | 0 | 5 |
| JC | 40.852 | -119.567 | 1443 | *hololeuca* | 0.000 | 0 | 0 | 15 | 15 |

| | | | | | | | | | |
|---|---|---|---|---|---|---|---|---|---|
| JO | 36.995 | -111.595 | 1249 | *juncea* | 0.894 | 5 | 0 | 0 | 5 |
| JT | 35.905 | -111.548 | 1504 | *juncea* | 0.889 | 6 | 0 | 0 | 6 |
| LA | 35.206 | -106.497 | 1906 | *latisquamea* | 0.384 | 0 | 6 | 0 | 6 |
| LO | 35.033 | -110.645 | 1487 | *bigelovii* | 0.994 | 5 | 0 | 0 | 5 |
| LR | 42.108 | -119.741 | 1467 | *hololeuca* | 0.011 | 0 | 0 | 7 | 7 |
| LT | 34.994 | -107.301 | 815 | *bigelovii* | 0.981 | 3 | 0 | 0 | 3 |
| LV | 39.626 | -120.010 | 1668 | *hololeuca* | 0.068 | 0 | 3 | 12 | 15 |
| MC | 45.999 | -110.833 | 1660 | unknown | 0.940 | 7 | 0 | 0 | 7 |
| M D | 41.411 | -120.544 | 1333 | *hololeuca* | 0.010 | 0 | 0 | 15 | 15 |
| MR | 45.999 | -110.833 | 1663 | unknown | 0.936 | 2 | 0 | 0 | 2 |
| NH | 38.024 | -117.869 | 1440 | *mohavensis* | 0.759 | 0 | 5 | 0 | 5 |
| NI | 35.958 | -111.767 | 1935 | *nitida* | 1.000 | 6 | 0 | 0 | 6 |
| NN | 41.073 | -115.292 | 1633 | unknown | 0.626 | 5 | 0 | 3 | 8 |
| NO | 39.226 | -114.903 | 2035 | *speciosa* | 0.247 | 0 | 5 | 0 | 5 |
| NT | 38.168 | -116.439 | 1910 | *speciosa* | 0.142 | 0 | 5 | 0 | 5 |
| NV | 40.595 | -116.173 | 1476 | *hololeuca* | 0.338 | 2 | 0 | 4 | 6 |
| OO | 38.578 | -112.334 | 1789 | *hololeuca* | 0.356 | 0 | 6 | 0 | 6 |
| OT | 38.717 | -112.375 | 1914 | *salicifolia* | 0.896 | 4 | 0 | 0 | 4 |
| PB | 43.930 | -121.106 | 1200 | unknown | 0.030 | 0 | 0 | 6 | 6 |
| PL | 39.607 | -119.899 | 1724 | *hololeuca* | 0.000 | 0 | 0 | 14 | 14 |
| PT | 39.505 | -119.902 | 1421 | *hololeuca* | 0.224 | 0 | 7 | 7 | 14 |
| PW | 38.595 | -118.115 | 1859 | *hololeuca* | 0.066 | 0 | 3 | 8 | 11 |
| RH | 45.729 | -119.047 | 318 | *speciosa* | 0.370 | 0 | 7 | 0 | 7 |
| RL | 43.529 | -119.299 | 1390 | unknown | 0.015 | 0 | 0 | 6 | 6 |
| RO | 44.147 | -117.107 | 881 | *speciosa* | 0.145 | 0 | 6 | 0 | 6 |
| RS | 37.515 | -113.211 | 1628 | *graveolens* | 1.000 | 6 | 0 | 0 | 6 |
| RT | 45.345 | -118.227 | 883 | *oreophila* | 0.287 | 0 | 2 | 0 | 2 |
| SC | 39.383 | -117.624 | 1990 | *hololeuca* | 0.002 | 0 | 0 | 14 | 14 |
| SH | 41.324 | -117.706 | 1487 | unknown | 0.012 | 0 | 0 | 5 | 5 |
| ST | 40.237 | -109.085 | 1727 | *turbinata* | 1.000 | 6 | 0 | 0 | 6 |
| SJ | 40.237 | -109.085 | 1727 | unknown | 0.999 | 6 | 0 | 0 | 6 |
| SL | 35.219 | -111.642 | 2140 | unknown | 1.000 | 5 | 0 | 0 | 5 |
| SS | 40.737 | -120.317 | 1647 | *hololeuca* | 0.135 | 0 | 4 | 10 | 14 |
| TO | 37.283 | -113.306 | 1159 | *salicifolia* | 0.239 | 0 | 6 | 0 | 6 |
| TU | 37.226 | -113.378 | 1004 | unknown | 1.000 | 7 | 0 | 0 | 7 |
| UN | 37.116 | -111.977 | 1447 | unknown | 0.292 | 0 | 2 | 0 | 2 |
| UO | 39.085 | -113.510 | 1438 | *speciosa* | 0.203 | 0 | 4 | 0 | 4 |
| UT | 39.073 | -113.720 | 1671 | *turbinata* | 1.000 | 5 | 0 | 0 | 5 |
| VI | 34.590 | -117.259 | 869 | *speciosa* | 0.164 | 0 | 5 | 0 | 5 |

| | | | | | | | | | |
|---|---|---|---|---|---|---|---|---|---|
| VM | 39.915 | -119.900 | 1503 | *hololeuca* | 0.000 | 0 | 0 | 15 | 15 |
| WA | 43.825 | -117.641 | 762 | *oreophila* | 0.940 | 3 | 0 | 0 | 3 |
| YL | 43.539 | -119.437 | 1307 | *speciosa* | 0.015 | 0 | 0 | 5 | 5 |

**Table S2: Deviance information criterion (DIC) values for different ancestral clusters (*k*) outputted from ENTROPY.** *k* = 2 model fit the data best indicated by lowest DIC value and presented in bold.

| *k* | chain 1 | chain 2 | chain 3 | chain 4 | mean |
|---|---|---|---|---|---|
| **2** | 31235761.82 | 31363445.98 | 31660942.42 | 31825995.42 | **31518422.34** |
| 3 | 33145796.79 | 33228694.28 | 33739298.47 | 34535151.69 | 33602822.33 |
| 4 | 34280800.92 | 34847580.16 | 35433797.03 | 35667550.29 | 35085184.27 |
| 5 | 33518588.06 | 33926279.57 | 34256212.05 | 34466287.03 | 34058309.72 |
| 6 | 35658707.76 | 36304858.22 | 37514473.47 | 37572699.02 | 36811678.36 |
| 7 | 36777341.52 | 36976124.13 | 38706378.32 | 41239748.01 | 38230349.07 |
| 8 | 83617352.63 | 133895982.7 | 235957096.2 | 301829319.8 | 187525471.7 |
| 9 | 145255251.6 | 350119378 | 648345751.7 | 856240278.3 | 499737631.6 |

**Table S3: Environmental variable abbreviations with descriptions and units used in the text and figures, and the range of values for each variable.** Data for these variables are represented more fully in Table S4.

| Env. variable | Description (units) | Variable range |
|---|---|---|
| Elevation | Elevation (m) | 242 - 2443 |
| AnnPpt | Annual precip. (mm) | 112.97 - 582.27 |
| AnnTemp | Annual temp. (°C) | 3.89 - 17.58 |
| Slope | Slope (degrees) | 0 - 29.77 |
| Aspect | Aspect (degrees from North) | 0 - 180 |
| SoilAWC | Max. soil water capacity (cm) | 0.08 - 0.22 |
| Heatload | Heat load index[2] | 0.63 - 1.03 |
| CumlAET | Cuml. actual evapotranspiration (mm per year) | 105.4 - 352.41 |
| FallAET | Fall actual evapotranspiration (mm per Sep-Nov) | 1.28 - 48.27 |
| PrcpSeas | Precip. seasonality[1] | 0.1 - 0.93 |

1. PrcpSeas classes designated by Walsh & Lawler, 1981
2. HL calculated from McCune & Keon, 2002
